# Cheese3D: Sensitive Detection and Analysis of Whole-Face Movement in Mice

**DOI:** 10.1101/2024.05.07.593051

**Authors:** Kyle Daruwalla, Irene Nozal Martin, Linghua Zhang, Diana Naglič, Andrew Frankel, Catherine Rasgaitis, Xinyan Zhang, Zainab Ahmad, Jeremy C. Borniger, Xun Helen Hou

## Abstract

Facial expressions and movements, from a subtle and ephemeral grimace, to vigorous and rapid chewing, offer direct insights into the moment-to-moment change of neural and physiological processes. Mice, with discernible facial responses and evolutionarily conserved mammalian facial movement control circuits, provide an ideal model to unravel the link between facial movement and underlying physiological states. However, existing frameworks lack the spatial or temporal resolution to sensitively track all movements of the mouse face, due to its small and conical form factor. We introduce Cheese3D, a computer vision system that captures high-speed 3D motion of the entire mouse face (including ears, eyes, whisker pad, and jaw, while covering both sides of the face) using a calibrated six-camera array. The interpretable framework extracts dynamics of anatomically-meaningful 3D facial features in absolute world units at sub-millimeter precision. The precise face-wide motion data generated by Cheese3D provides clear physiological insights, as shown by proof-of-principle experiments predicting time under general anesthesia from changing facial patterns, inferring tooth and muscle anatomy from fast ingestion motions across the entire face, measuring minute differences in movements evoked by brainstem stimulation, and relating neural activity to spontaneous facial movements, including expressive features only measurable in 3D (e.g., angles of ear motion). Cheese3D can serve as a discovery tool that renders subtle mouse facial movements highly interpretable as a readout of otherwise hidden internal states.

## 1 Introduction

Facial expressions and movements, from grimacing to chewing, are a powerful reflection of our internal states in health and disease [1, 2]. Studying how coordinated motion of individual facial regions gives rise to multi-functional whole-face movements can therefore provide unique insights into internal physiological processes [3]. Work to-date suggests we can infer pain, distress, and sensory input based on subtle facial movement patterns in humans as well as rodents [4–12]. Thus, once deciphered, the face can serve as a high-bandwidth and compact window into the unseen internal states of animals. Mice share evolutionarily conserved facial movement control circuits with other mammals, including humans. Facial muscles controlling eyes, ears, whiskers, nose, and mouth receive direct commands from a motor control network in the brainstem, bypassing the spinal cord, and thus, are positioned relatively close to processing centers in the brain [13–15]. This shared circuit architecture makes laboratory mice ideally suited to serve as a model system for studying the link between facial movements and internal brain and body states.

Realizing the potential of face as an informative readout requires a framework with the sensitivity, precision, and accuracy to quantitatively relate facial movement to internal state. Although recent advances in computer vision have fueled state-of-the-art methods for human facial movement tracking [16, 17], similar approaches to characterizing face-wide movements in mice encounter unique technical challenges. Mouse faces are an order of magnitude smaller than human faces, and the conical shape of their head makes it difficult to capture face-wide movement using a single camera (**Figure 1a**). Existing methods rely on zooming into motion of a single facial region (e.g. whiskers, tongue) or a subset of facial regions on one side of the face [3, 18, 19]. Alternative methods have discarded temporal dynamics by focusing on still images of the face [6, 20]. Recent 3D methods hold promise to capture movements of the whole animal [21–23], but the approach has not been evaluated at the high-resolution required to examine the face of mouse.

**Figure 1:**
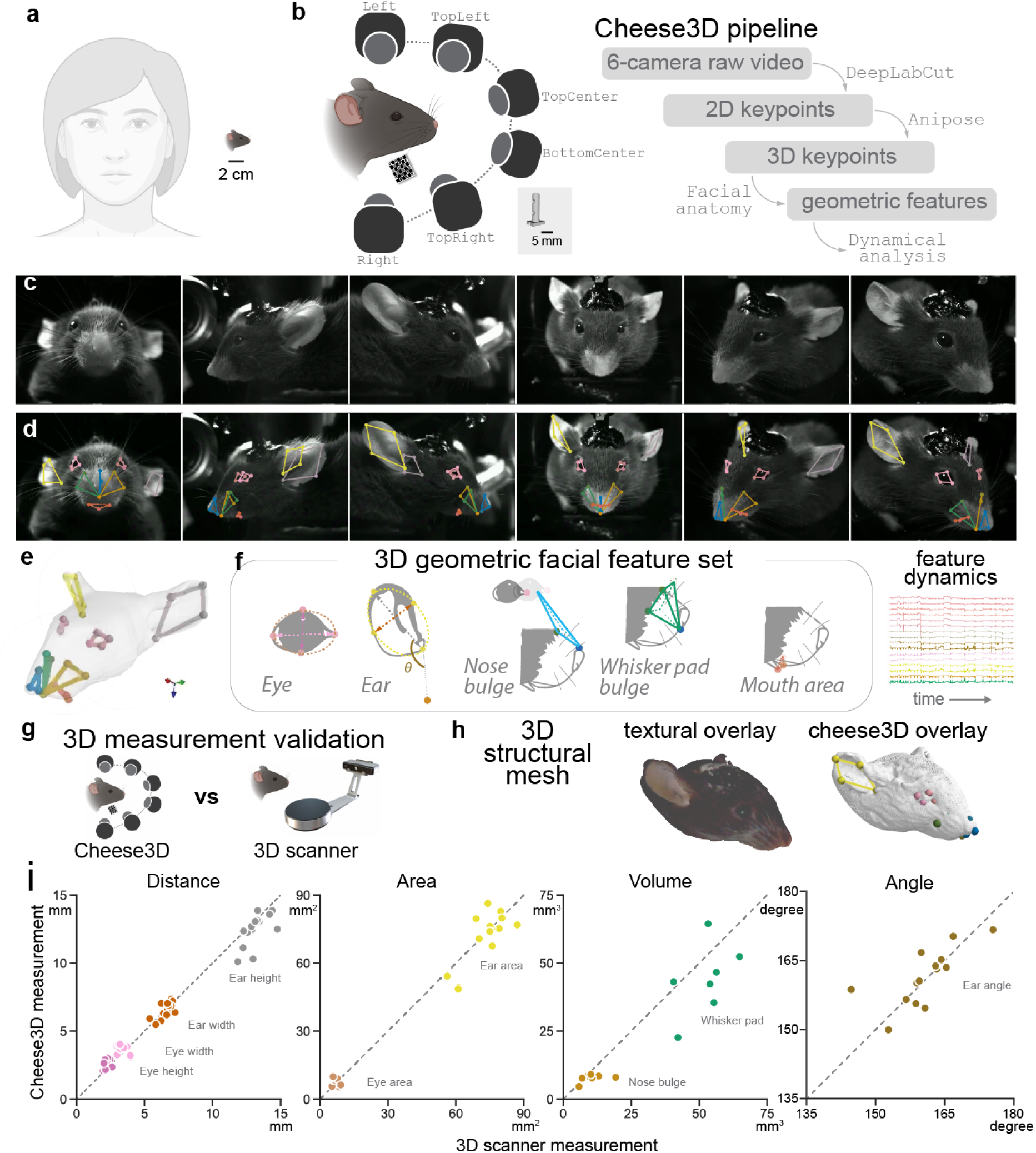
Framework and validation of capturing face-wide movement as 3D geometric features in mice **(a)** The form factor of mouse face poses technical challenges to track mouse face-wide movement, compared to existing technology tailored for the human face. **(b)** Schematic of the hardware and software framework. *Left*: The six-camera facial movement capture setup. The ChArUco board shown below the mouse is required for camera calibration. *Inset*: a head-post designed to image mouse face without occluding any facial features. *Right*: the analysis pipeline which inputs six-camera raw video and outputs the dynamics of a geometric facial feature set. **(c)** Example synchronized frames from the six-camera setup. **(d)** 3D facial keypoints visualized as projections onto the frames shown in **(c)**. **(e)** 3D facial keypoints overlaid onto a 3D template mouse face from [27] used purely as a visual aid. **(f)** Output of Cheese3D. *Left*: Illustrations of the set of anatomically-based facial features including 3D distances, areas, volumes, and angles across facial regions (see text and methods for details). *Right*, example time series of the 3D feature set. **(g)** Experimental design to validate Cheese3D facial feature measurement (anesthetized) compared to 3D scanner. **(h)** Example mouse face 3D mesh obtained via 3D scanner *Left*: with texture overlay, showing fur and color details. *Right*: the same mesh overlaid with 3D keypoints obtained from Cheese3D to compare the two. **(i)** Comparison of Cheese3D facial feature measurement with 3D scanner data, grouped by distances, areas, volumes, and angle, from left to right. Measurements for lateralized facial features (eyes, ears) contain both left and right sides and thus have twice the amount of data points compared to midline features (nose bulge, whisker pad bulge). Mouth area is the only feature excluded from the comparison since it cannot be reliably measured on the 3D scanner due to the orientation of the mouse face relative to the projector.

Adapting components from existing markerless pose estimation tools, we carefully designed a hardware and software pipeline to create a unified 3D view of the mouse face, and introduced technical advancements to reduce keypoint jitters common to existing tools, a critical step to increasing resolution and sensitivity necessary to study subtle and rapid mouse facial movements. Our approach, Cheese3D, balances the trade-offs in current tools, resulting in the first method sensitive enough to characterize how internal physiological, cognitive, and emotional states drive overt dynamics of the face. Cheese3D captures the whole face—including insightful features such as the ears and both sides of the face, absent in existing tools due to technical challenges—and a wide dynamic range of movements, revealing the potential of facial movement as a non-invasive, information dense readout of internal states and processes.

## 2 Results

### 2.1 Cheese3D captures robust 3D whole-face movement in mice

The Cheese3D pipeline captures and analyzes synchronous movement of the entire mouse face at 100 Hz temporal resolution. The three pairs of high-speed video cameras (six total) are positioned compactly to capture the frontal view (Top Center and Bottom Center cameras), the profile view (Left and Right cameras), and an elevated half-profile view (Top Left and Top Right cameras) (**Figure 1b**). To acquire high-resolution facial video while maintaining comfort for natural behavior, mice are acclimated to sitting in a tunnel with the head secured using a lightweight headpost custom-designed to allow unobstructed viewing of all facial areas (**Figure 1b** *inset*). Individual views from the six-camera array are temporally synchronized, and spatial alignment between views is captured through ChArUco calibration [24]. We identified a set of 27 facial keypoints that covers all facial areas on C57BL/6J mouse (**Figures 1c**–**1e**, **Supplementary Video 1**). Each keypoint is in sharp focus and visible by at least two cameras (see **Supplementary Table 1**), and reproducibly labeled by different researchers following written guidelines. The calibrated hardware setup and labeling protocol enables us to adapt existing markerless pose estimation techniques, such as Anipose [22] and DeepLabCut [23], to create a unified 3D view of the whole mouse face at the spatial and temporal resolutions necessary to study facial movements.

As facial movement is inherently constrained in 3D space, existing 2D methods for mouse facial analysis either require a single camera view, limiting the type of movement studied to those that can be captured in a single plane, or relies on principal component analysis or hidden Markov models to integrate keypoints across multiple uncalibrated views, hindering direct interpretation [3, 25]. Moving from 2D to 3D calibrated face space is critical to enable interpretable feature selection that is physically grounded and verifiable in world units: we selected a set of 17 3D geometric features—distances, angles, areas, and volumes in 3D space—constructed from shapes defined by facial keypoints (**Figure 1f**). Furthermore, the features are localized to facial regions based upon known muscular anatomy and descriptors of rodent facial movements [5, 26].

We evaluated the accuracy of Cheese3D by comparing its resulting 3D geometric features with those measured statically using a 3D scanner (resolution: 50 µm) for the same mouse (**Figures 1g**–**1i**, **Supplementary Figure 1, Supplementary Video 2**). (Mean *±* RMSE. Eye height: 2.62 *±* 0.52 mm; Eye width: 3.71 *±* 0.63 mm; Ear height: 12.45 *±* 1.13 mm; Ear width: 6.54*±*0.43 mm; Ear angle: 161.45*±*4.86*^◦^*; Eye area: 8.14*±*2.27 mm^2^; Ear area: 71.18*±*7.39 mm^2^; Nose bulge: 7.74 *±* 4.75 mm^3^; Whisker pad bulge: 43.87 *±* 13.57 mm^3^; *n* = 7 mice) (**Figure 1i**).

To validate the utility and necessity of having all six cameras, we omitted different pairs of cameras and measured changes in accuracy in corresponding facial regions (**Supplementary Figure 2**). Omitting frontal cameras resulted in skewed measurements of midline facial features (e.g. whisker pad bulge), whereas omitting elevated half-profile cameras resulted in errors of the most lateral features (e.g. ear). The six-camera array is also essential as it builds in redundancy which ensures measurements are still possible even when part of the face is obstructed in some views, as is often the case when the mouse paws (e.g. during grooming) or experimental apparatuses (e.g. to deliver food, drugs, or olfactory stimuli) come into close proximity to the face. Collectively, the synchronized and calibrated array of six cameras, combined with geometric facial features in 3D, reduces the tradeoff between compromising spatial versus temporal resolution in characterizing rodent facial movement.

**Figure 2:**
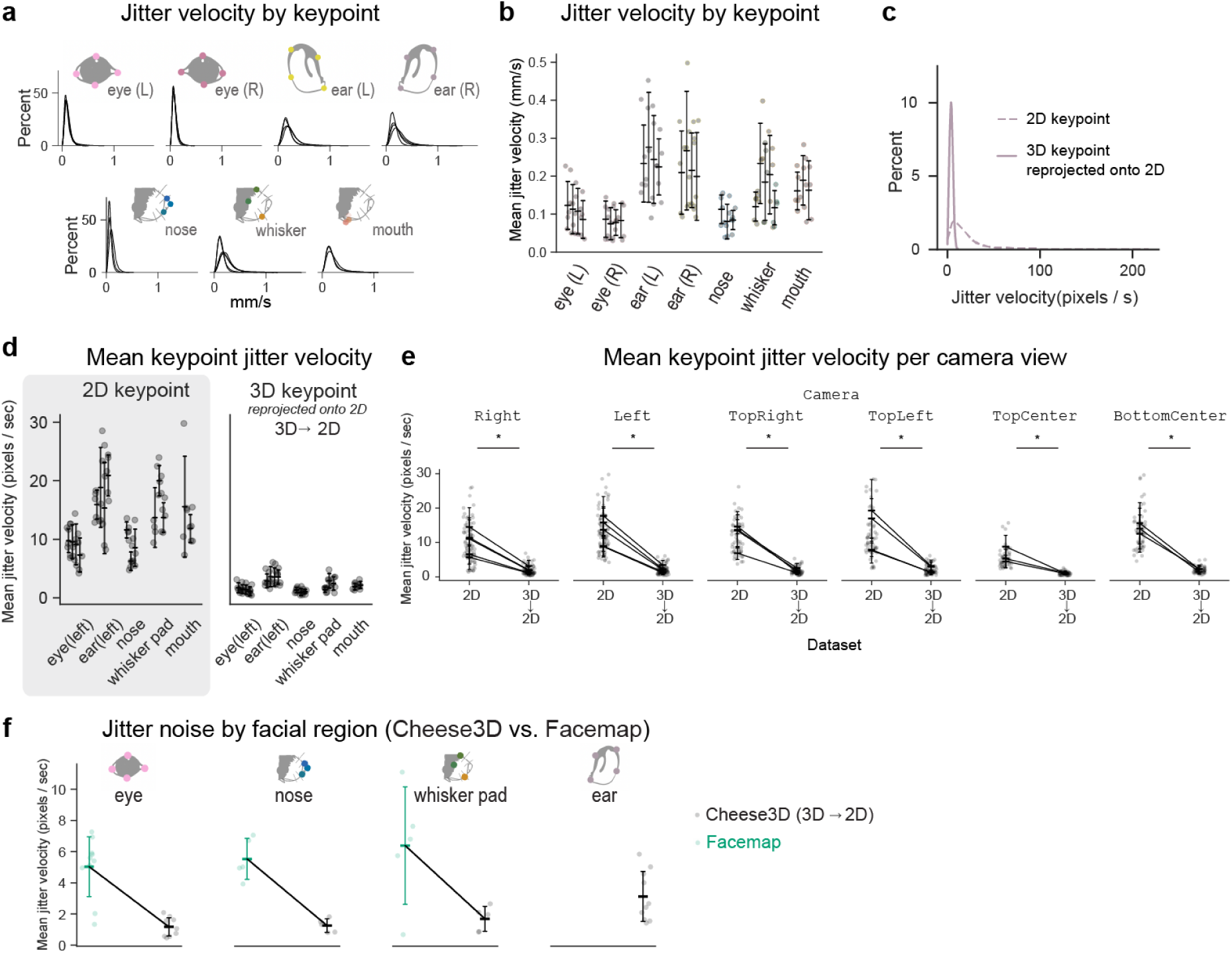
Reduction in keypoint tracking jitter due to 3D triangulation of data from six camera views **(a)** Distribution of keypoint-specific jitter (frame-to-frame velocity) during a motionless period for an example mouse. Each subpanel indicates a different facial region, and each curve indicates a different keypoint. **(b)** Summary of keypoint-specific jitter across mice (*n* = 5), where each column group indicates a facial region and each column indicates a keypoint. **(c)** Reduction of keypoint-specific jitter for an example keypoint from the left ear region in left camera view for an example mouse in 2D keypoints (prior to triangulation, dashed line) compared to 3D keypoints reprojected onto the 2D camera view plane (after triangulation, solid line). **(d)** Changes in jitter for all keypoints from one example view (left camera) across mice (*n* = 5) between pre-triangulation (2D keypoints, shaded region) and post-triangulation (reprojected 2D keypoints). See **Supplementary Figure 3** for all views and summary statistics. **(e)** Summary of keypoint-specific jitter across mice and all six camera views. See **Supplementary Figure 3** for summary statistics. **(f)** Jitter comparison between Cheese3D (black) and Facemap (green) across facial regions. Each point represents the mean jitter velocity across all keypoints for the facial regions as well as camera views for a single mouse. Cheese3D’s 3D keypoints are reprojected onto 2D views to compare with Facemap. Note Facemap’s output does not include the mouse’s ears. Measurements for lateralized facial features (eyes, ears) contain both left and right sides and thus have twice the amount of data points compared to midline features (nose, whisker pad). See **Supplementary Figure 5** for the same data broken down by camera view.

### 2.2 Reduction of tracking noise enables precise measurement of subtle and transient movements across facial regions

To assess the sensitivity of Cheese3D to detect and measure small, localized facial movements, we sought to explicitly quantify keypoint jitter in our setup in a control experiment using motionless periods during anesthesia. Keypoint jitter is a known issue whereby local fluctuations in keypoint tracking are unrelated to genuine movement [25]. This can happen due to image noise, inadequate lighting, low contrast/texture, label noise in the training data, or keypoint-specific uncertainty in the model. Across the facial keypoints selected, human labelers utilize not only texture, but also color and shape to determine the location of keypoints. Convolutional neural networks often focus on texture to solve object recognition tasks [28], thus it should be expected that certain keypoints which rely primarily on texture for their location are learned more confidently than others. Keypoint jitters are often mitigated using low-pass filters, but this can attenuate dynamics and reduce temporal resolution of detection [25]. A critical benefit of 3D multi-view calibration compared to single 2D uncalibrated views is that view redundancy reduces the amplitude of keypoint jitter, allowing us to detect more subtle fast movements. Studying motionless periods, we detected jitters of 3D keypoints without any filtering (**Figures 2a**, **2b**. Mean *±* std, grouped by facial regions. Ear (left): 0.24 *±* 0.10 mm/sec; Ear (right): 0.22 *±* 0.11 mm/sec; Eye (left): 0.11 *±* 0.06 mm/sec; Eye (right): 0.08 *±* 0.04 mm/sec; Nose: 0.09 *±* 0.04 mm/sec; Whisker pad: 0.17 *±* 0.09 mm/sec; Mouth: 0.17 *±* 0.06 mm/sec; *n* = 5 mice), and measured the reduction in jitter between 2D keypoints and 3D keypoints projected onto 2D views (**Figures 2c**–**2e**; all *p <* 0.05, one-sided Wilcoxon matched-pairs test, 2D keypoints *>* 3D keypoints reprojected onto 2D; see **Supplementary Figure 3** for more details). Additionally, we processed the same video data using an existing 2D mouse face pose tracking tool, Facemap [3] (**Figure 2f**, **Supplementary Figure 5**. For Facemap: Eye: 5.04 *±* 1.92 px/sec, Nose: 5.53 *±* 1.32 px/sec, Whisker pad: 6.39 *±* 3.78 px/sec; For Cheese3D: Eye: 1.17 *±* 0.59 px/sec, Nose: 1.26 *±* 0.44 px/sec, Whisker pad: 1.69 *±* 0.80 px/sec, Ear: 3.12 *±* 1.61 px/sec; *p <* 0.005, one-sided Wilcoxon matched-pairs test (Facemap > post-triangulation Cheese3D)). In all comparisons, we observed a reduction in keypoint jitter after 3D triangulation. We further examined the effect of keypoint jitter on geometric features, which informs the mouse-specific threshold between keypoint tracking noise and bona fide movements that Cheese3D can detect (**Supplementary Figure 4**. 99.9th percentile of jitter noise: Ear angle (left): 5.01 *±* 2.39 °/sec; Ear angle (right): 4.50 *±* 2.22 °/sec; Eye area (left): 1.63 *±* 0.86 mm^2^/sec; Eye area (right): 1.24 *±* 0.66 mm^2^/sec; Mouth area: 0.88 *±* 0.41 mm^2^/sec; Whisker pad bulge: 7.71 *±* 2.33 mm^3^/sec; Nose bulge: 2.18 *±* 0.83 mm^3^/sec; *n* = 5 mice).

### 2.3 Uncovering underlying physiology from external facial movements

As a proof-of-principle test that Cheese3D is able to capture subtle and rapid facial movements with physiological significance, we designed experiments to monitor mice emerging from ketamine-induced general anesthesia. Small localized facial movements, including whisker deflections, both appear during anesthesia as well as signal early stages of recovery [29]. This poses unique challenges to sensitively track subtle movements with coverage of the entire face, compared to other overt body movements such as locomotion and reaching, where limbs and appendages undergo large translations and rotations relative to their size. We applied the thresholds defined by the jitter analysis (above in **Supplementary Figure 4c**) to facial movements recorded by Cheese3D during anesthesia, a common procedure that is associated with subtle facial movements [29]. A detailed examination of facial movement velocity during induction, maintenance, and recovery of anesthesia reveals temporal patterns across different facial regions. Motions within different facial regions are visualized in the movement raster plot, where each vertical line represents displacement above jitter threshold in the corresponding video frame (we selected one representative geometric feature per each of the seven facial regions) (**Figures 3a**, **3b**). These data demonstrate that Cheese3D can be used to detect small movements associated with anesthesia and recovery in mice.

**Figure 3:**
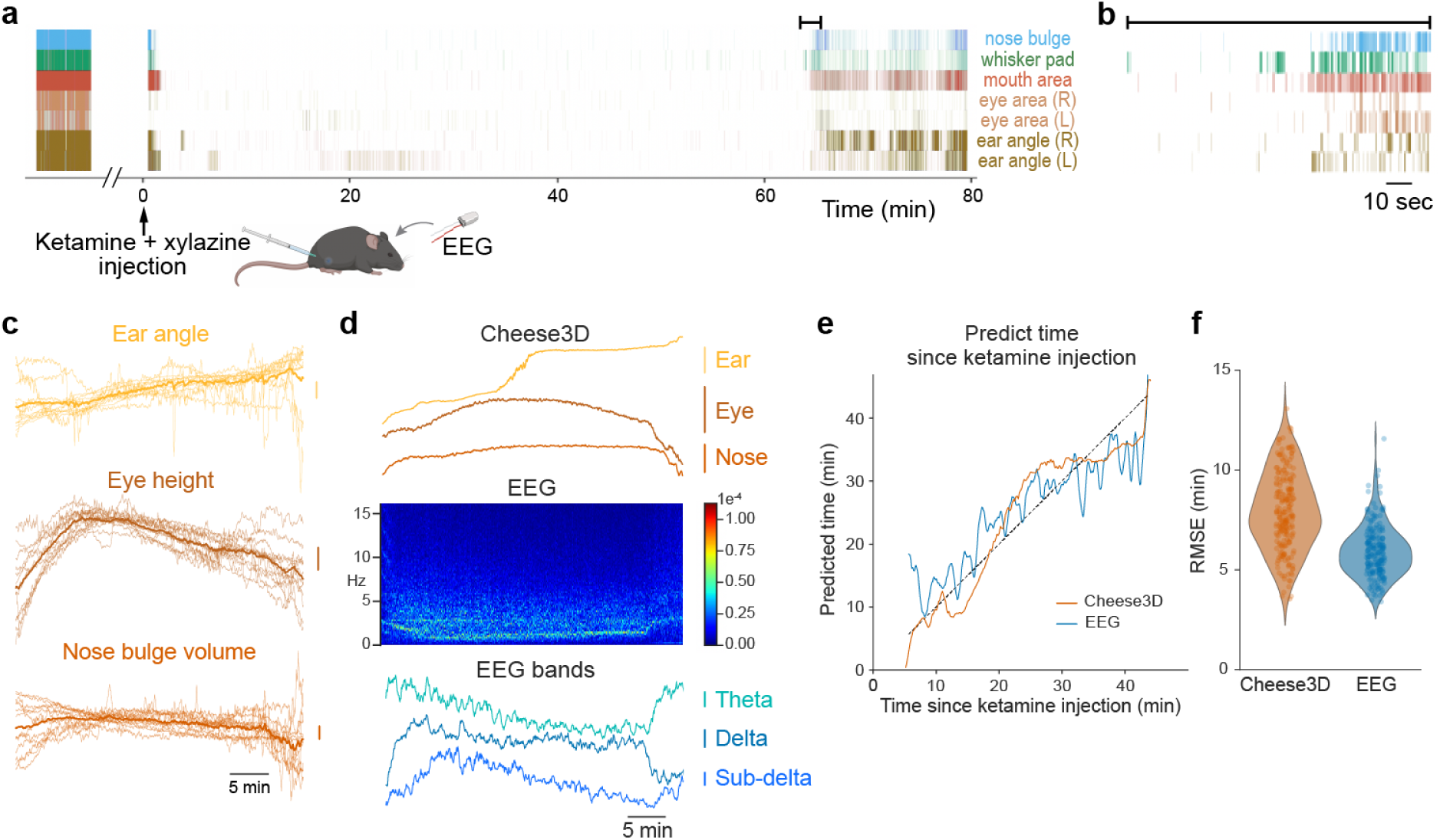
Distinct facial patterns track time during induction and recovery from ketamine-induced anesthesia **(a)** Example facial movement raster plot during anesthesia with concurrent EEG recording (each vertical line corresponds to movement above the 99.9-th percentile jitter threshold as shown in **Supplementary Figure 4c** for a given 10 ms time window). **(b)** Zoom in of movement raster plot from **(a)** to show the early moments of movement recovery following anesthesia. **(c)** Overlaid z-scored ear angle, eye height, and nose bulge volume traces from 12 sessions (*n* = 3 mice, four sessions per mouse. Vertical scale bar indicates one standard deviation). **(d)** Simultaneously recorded Cheese3D features (*top*, showing moving average over a 10 sec window; vertical scale bars: ear: 2*^◦^*, eye: 0.1 mm, nose: 0.5 mm^3^), EEG spectrogram (*middle*, 5 sec FFT window), and power of EEG frequency bands (*bottom*, showing subdelta: 0.2 Hz to 1 Hz, delta: 1 Hz to 4 Hz, and theta: 5 Hz to 10 Hz bands; vertical scale bars indicate one standard deviation) for an example session. **(e)** Output from a quadratic model fit across mice predicting time since injection using the initial and current Cheese3D (orange) or EEG (blue) feature values relative to the dotted identity line. **(f)** Root-mean-square error (RMSE) of time prediction where each dot represents the mean test error for one particular model trained on either Cheese3D (orange) or EEG (blue) features.

We examined if Cheese3D can be used to study facial movements associated with physiological processes that are not otherwise externally visible. Visualizing the anesthesia data over the entire period from induction to recovery revealed gradual changes in facial motion (e.g., ears and eyes) that are stereotyped across sessions and across mice (**Figure 3c**, **Supplementary Video 3**). This suggests that certain facial features can be used as a “stopwatch” across mice to track time since anesthesia induction. To test this hypothesis, we fit a single model across all mice to predict the time elapsed since anesthesia induction (intraperitoneal injection of ketamine and xylazine) using only the initial and current value of filtered (10 sec moving average window) facial features (**Figures 3e**, **3f**). As a comparison, we recorded temporally synchronized electroencephalography (EEG) and fit the same model using power from EEG frequency bands (**Figure 3d**). We observe that Cheese3D facial features allow us to predict time since anesthesia induction with similar performance to using EEG features (**Figure 3f**; RMSE: 7.91*±* 1.95 min for Cheese3D, 5.90*±* 1.27 min for EEG; *p* = 0.068, paired two-sided t-test with *k*-fold cross-validation correction [30, 31], 4 sessions per mouse, *n* = 3 mice). In short, combining motions from different facial regions can serve as a useful visual indicator of time elapsed since induction of general anesthesia.

We tested Cheese3D with facial movements that are vigorous in amplitude: chewing in rodents is difficult to characterize externally as teeth, along with food that has entered the mouth, cannot be seen. However, being able to track and measure chewing is essential to studies of nutrient absorption and efficient digestion [32] as well as jaw proprioception [33]. Existing techniques to characterize chewing rely on invasive methods such as electromyography, to infer what is happening inside the mouth [34]. We hypothesized that Cheese3D would enable more direct assessment of chewing dynamics from careful examination of external facial movements during food consumption. Using the same Cheese3D multi-camera array and 17 geometric facial feature identification system with no modifications, we recorded mice as they ate crunchy food (3 mm diameter precision pellets), and visualized 3D trajectory of mouth keypoints (upper lip corners and lower lip, forming a triangle in 3D space; **Figures 4a**, **4b**). Plotting the area of this triangle (i.e., mouth opening) over time revealed two distinct modes of eating, with either elevated or reduced lower signal envelope, corresponding respectively to a food pellet obstructing the mouth opening or the mouth shut (**Figure 4c**). The transition between the two modes is abrupt and reliably identifiable across all mice (5.20 *±* 1.71 sec; ranging from 2.77 sec to 8.12 sec; *n* = 7 mice. **Figure 4d**, **Supplementary Video 4**). The clear separation is also evident in movements within the facial area close to the back of the mouth (**Figures 4e**, **4f**). This finding is consistent with the unique tooth anatomy of rodents, in which a distinct gap, termed diastema, separates the incisors (for ingestion) from the molars (for mastication), as labeled in microCT images (**Figure 4g**). Whole-face movement analysis also revealed temporally correlated eye protrusion with chewing during mastication but not ingestion or other spontaneous facial movement for every mouse examined (peak cross-correlation: 38.05 *±* 19.45 for mastication, 2.80 *±* 2.81 for ingestion; *n* = 7 mice; *p <* 0.001, one-sided Wilcoxon matched-pairs test (mastication mean value *>* ingestion mean value); **Figures 4h**–**4k**, **Supplementary Video 5**). This could potentially be attributed to the anatomy of the rodent muscles of mastication since they wrap around the base of the eye socket [26]. The phenomenon has been frequently observed and named “eye boggling” in the pet rodent community, and only recently described in scientific literature [35]. Our data indicate that Cheese3D detects facial movements during rodent food consumption consistent with known characteristics of food placement, tooth anatomy, and muscle engagement.

**Figure 4:**
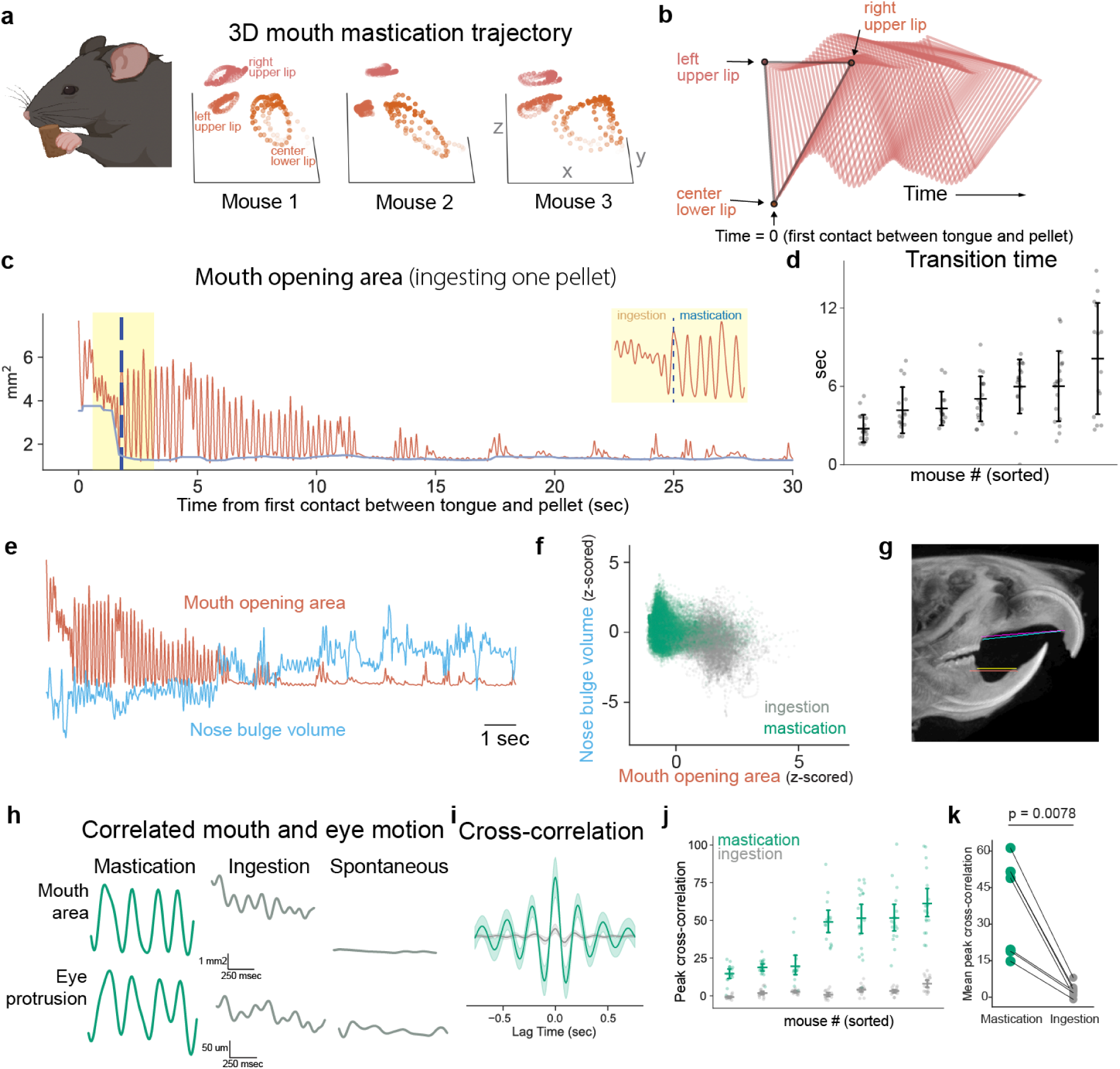
Chewing kinematics in mouth and surrounding facial areas **(a)** 3D trajectory of three mouth keypoints for 1 sec of chewing motion for three example mice. **(b)** Time evolution of the mouth opening triangle formed by the three mouth keypoints in **(a)** for 0.5 sec from the moment the food pellet comes into contact with the tongue for an example mouse. **(c)** Area of the mouth opening triangle over time during consumption of one pellet for an example mouse. Blue dashed vertical line indicates putative transition time from ingestion (incisor chewing) to mastication (molar chewing), with a zoomed in view around the transition time (yellow shaded area) shown in *inset*. **(d)** Summary of ingestion to mastication transition times. Each column is one mouse; each dot is one food pellet. **(e)** Visualizing mouth opening area concurrent with nose bulge volume (z-scored per feature) while an example mouse consumes a single pellet (same pellet as shown in **(c)**). **(f)** Same data as in **(e)** where each dot represents a time point, colored based on before the transition time (gray, putative ingestion phase) or after (green, putative mastication phase). **(g)** MicroCT image of the mouse with diastema, the gap between incisors (for ingestion) and molars (for mastication), labeled in color lines. **(h)** Example time segments of mouth area with eye protrusion, during putative mastication (green), ingestion (gray), and during spontaneous movement outside of chewing (gray). **(i)** Cross-correlation between mouth area and eye protrusion for one example session for putative mastication (green) and ingestion (gray) phases. **(j)** Summary of peak cross-correlation (computed as shown in **(i)**) across pellets, where each column is one mouse. **(k)** Summary of mean peak cross-correlation (computed as shown in **(h)**) across pellets, where each point is one mouse (*n* = 7 mice).

### 2.4 Linking facial movement to motor control machineries using synchronized Cheese3D with electrophysiology

Facial musculature is controlled by motor neurons originating from the brainstem motor network. As such, relating facial movement to facial motor control machineries is a key first step to delineating the mechanisms linking brain-wide activity to facial behavior. Thus far, we have demonstrated that Cheese3D is accurate and precise enough to reveal internal physiological processes such as anesthetic depth and food ingestion phases. Yet, facial movements are unique in that large internal differences can manifest as subtle changes between two nearly identical movements. To quantify whether Cheese3D has the precision to detect such distinctions, we designed an experiment to induce small, localized movements in anesthetized mice via focal electrical stimulation of the brainstem motor control network using a four-shank silicon probe. By varying the current amplitude and location of stimulation, we aimed to create minute differences in facial responses engaging the building blocks of facial movement across different regions (**Figure 5a**).

**Figure 5:**
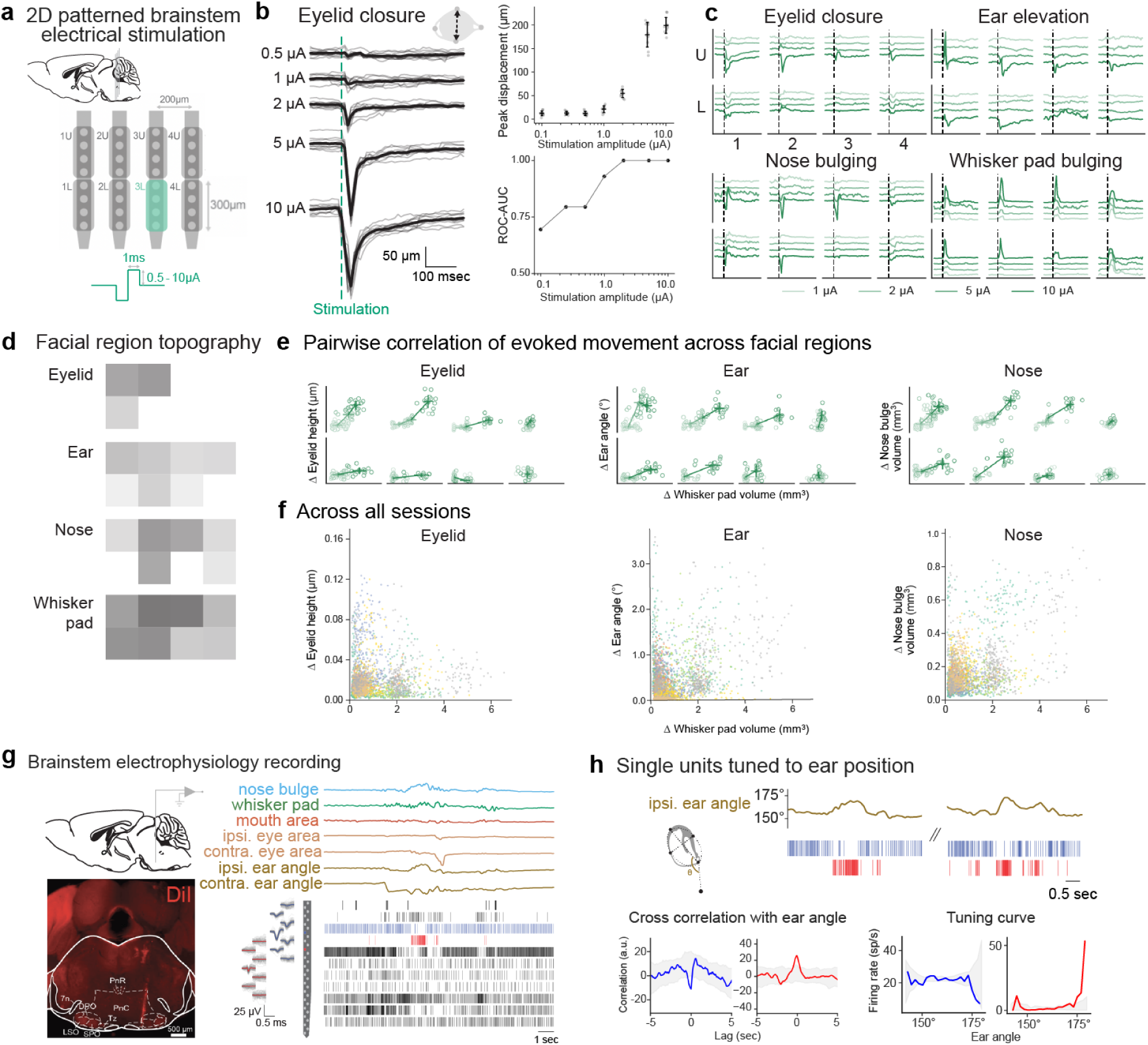
Synchronized Cheese3D with electrophysiology relates motor control activity to subtle facial movements **(a)** Schematic showing brainstem targeted by four-shank silicon probe. Electrodes are divided into a two by four grid to allow for patterned stimulation of the targeted area. **(b)** Example traces of ipsilateral eyelid closing triggered by bulk electrical stimulation across varying current amplitudes (*n* = 10 stimulations per amplitude, left) with peak absolute deflection measured as a function of stimulation amplitude (top right). Area under receiver operating characteristic curve when distinguishing pre-stimulation responses from post-stimulation responses for varying stimulation amplitudes (bottom right). **(c)** Responses across facial features to spacial-specific focal stimulation at varying amplitudes. Each trace indicates the mean response per amplitude (*n* = 15-30 stimulations per amplitude and location). **(d)** Peak absolute deflection of responses in **(c)** at 10 µA showing spatially localized movement for different facial features. **(e)** Pairwise relationship of peak absolute deflections of responses in **(c)** relative to whisker pad volume. See **Supplementary Table 2** for summary statistics. **(f)** Pairwise relationship of peak absolute deflections of responses as in **(e)** but colored by experimental sessions (9 sessions from *n* = 4 mice). **(g)** Schematics of in vivo electrophysiological recordings (top left) with silicon probe track revealed by DiI in the brainstem PnC (bottom left). Example recording of facial movements and single-unit activity in the brainstem of awake mice (5 sessions from *n* = 3 mice, 42 units, right). PnC: pontine reticular nucleus caudalis. PnR: pontine raphe nucleus. DPO: dorsal periolivary region. LSO: lateral superior olive. SPO: superior paraolivary nucleus. Tz: nucleus of the trapezoid body. 7n: facial nerve. **(h)** Two example single units with anti-correlated activities that are both tuned to ear angle (same as the blue and red units in the spike raster in **(g)**. Neural activities (top), cross correlation with ipsilateral ear angle (bottom left), and tuning curves (right) for the two highlighted example units in **(g)**. The lag of peak correlation is *−*20 ms and *−*80 ms, respectively. Gray shading in cross correlation: 99.9% confidence interval (CI) of shuffled data. Gray shading in tuning curve: 95% CI of shuffled data.

What is the smallest facial movement Cheese3D can reliably detect? First, to ensure that Cheese3D can detect controlled differences in facial movement, we bulk stimulated the entire region targeted by our probe at varying current amplitudes. We observed deflections in various facial features such as the distance between eyelids (called “eye height” in our tool) (**Figure 5b**; Mean *±* std by amplitude: 0.1 µA: 12.32 *±* 5.10 µm, 0.25 µA: 12.80 *±* 3.70 µm, 0.5 µA: 12.07 *±* 4.52 µm, 1.0 µA: 20.99 *±* 7.06 µm, 2.0 µA: 54.55 *±* 8.69 µm, 5.0 µA: 179.27 *±* 26.13 µm, 10 µA: 199.01 *±* 16.84 µm; *n* = 10 stimulations per amplitude). Notably, we detected changes in the eye height, as small as 2.66 µm, that increased with stimulation current, suggesting that Cheese3D can measure subtle differences in facial movement caused by neuronal activity. Our ability to distinguish such movements from pre-stimulation null responses improves with stimulation current amplitude (**Figure 5b**, bottom right). Next, we applied focal stimulation at varying current amplitudes in an effort to target the building blocks of facial movement in the brainstem. While we continued to observe current amplitude dependent changes, we also observed spatially localized movement responses for different facial features (**Figures 5c**, **5d**). Cheese3D facial features include both highly local features (e.g., eye height) that relate to localized musculature (e.g. eyelid closure) [36]) and global features (e.g., whisker pad volume) that cover a larger region of the face and are affected by the activity of multiple muscles. For these global features, we see movements due to stimulation for all locations, but the modulation of responses is location specific (**Figures 5c**, **5d**). This highlights the ability of global features to quantify complex whole-face motion. For example, when we examined the pairwise movement responses of various features against the whisker pad volume, we observed that locations with strong eye movement and whisker pad movement responses group into distinct current amplitude dependent clusters (**Figure 5e**). On the other hand, for regions with little eye movement, clusters do not form even though there are measurable deflections in the whisker pad volume. Across experimental sessions, we observe solely local feature responses as well as a mix of global and local facial responses (**Figure 5f**). Thus, while local features describe the movement of a single facial region, global features summarize how these parts work in synchrony across the face. In addition to local vs. global responses, we also observed asymmetry across the ipsilateral and contralateral sides of the face (**Supplementary Figure 7**). These complex motions indirectly reflect underlying biological mechanisms for coordination, but, as stimulation-evoked motion has demonstrated, Cheese3D has the precision to be one piece of a comprehensive quantification of the building blocks of facial movement.

Electrophysiological recordings in awake mice synchronized with Cheese3D further demonstrated correlated brainstem neural activity with specific facial features during spontaneous movements (**Figure 5g**). Measuring facial motion in calibrated 3D world coordinates enables detecting expressive facial features with complex movements in 3D space, such as elevation and abduction of the pinna (3D “ear angle” in Cheese3D output). Indeed, we discovered single units in the pontine reticular nucleus caudalis (PnC), tuned to ear movement, with firing rates adjusting to direction-specific ear movements (**Figures 5g**, **5h**, in blue and red; *p <* 0.001), as suggested by prior work [37, 38]. With quantitative data from Cheese3D, we can precisely calculate the units’ tuning curves with respect to ear position.

Taken together, Cheese3D has the precision and accuracy to relate the activity of neurons in the brainstem to facial movements. We demonstrated this by both inducing movements through directed stimulation, as well as relating recorded neural activity to spontaneous motion. In particular, we demonstrated the ability to measure small differences in nearly identical motions, illustrating Cheese3D’s ability to capture the subtleties of facial movements. Thus, Cheese3D is a powerful tool for uncovering the mechanism underpinning facial motion.

## 3 Discussion

The goal of Cheese3D is to provide an interpretable framework for using mouse face-wide movement to discover underlying physiological functions across a wide range of applications. Recognizing the unique and underexplored potential to use whole-face dynamics as a noninvasive readout of moment-to-moment changes of body and brain states in mice, we crafted Cheese3D as a specialized high-resolution tool to study mouse facial movements, compared to and built upon emerging animal behavioral tracking methods aimed to generalize across body parts and species [21–23, 39–44]. Moreover, in contrast to existing methods that focus on static facial images, motion of a subset of facial features, or aggregates of orofacial behavior optimized to predict cortical neural activities [3, 6], Cheese3D is specifically designed to capture and represent whole-face movement while maintaining spatial and physical interpretability. The multi-camera array setup facilitates reliable, markerless identification of facial keypoints in 3D space, counteracting occlusion and distortion found in single-camera setups, and reduces keypoint jitter compared to 2D methods. These precise spatial locations relative to other facial regions are preserved in the 3D geometric features. Recording the motion of both individual facial regions and their spatial and temporal relationship to the whole face is meaningful, since the building blocks of facial movement, i.e., compartments of facial musculature and the brainstem nuclei that directly control them, are highly topographically arranged [13, 14].

The proof-of-principle work demonstrates the utility of Cheese3D in detecting and characterizing both subtle movements as well as significant and temporally variable movements. Our analysis revealed informative synchronous facial movement patterns that could be used to infer unseen (internal anatomy and physiological functions) from seen (external synchronized facial motion). In particular, subtle differences between two nearly identical facial movements have the capacity to reflect large internal changes. Through electrical stimulation in brainstem, we demonstrate Cheese3D’s ability to detect such differences, illustrating its ability to reveal the potential of facial behavior as an informative, high-bandwidth readout of internal state. Moreover, spatially varied facial responses revealed by patterned stimulation in brainstem suggests Cheese3D, coupled with precisely synchronized electrophysiology, can tap into the functional organization of facial control machineries, adding to existing knowledge inferred from anatomical tracing. Indeed, our preliminary findings in brainstem recordings of awake mice identified potential PnC neurons correlating with the directional ear movement. Although not in the scope of the current work, the framework described in detail here can be adapted for different strains of mice, in freely moving setup, as well as for tracking development. We anticipate the method will enable important discoveries across fields in biology and medicine by allowing for noninvasive readout of moment-to-moment changes in body states in mice. The potential applications of high-resolution, whole-face kinematics data made possible by Cheese3D are vast and are likely to inspire a new era of quantitative studies linking facial movements to changes in internal states brought on by disease, drug exposure, neural processes, or other physiological functions we would otherwise have limited access to based on external observations.

## 4 Methods

### 4.1 Mouse

All experiments were performed in compliance with protocols approved by the Institutional Animal Care and Use Committee at Cold Spring Harbor Laboratory (protocol number 22-6). Both female and male C57BL/6J mice 2-8 months of age were used for experiments. Unless stated otherwise, animals were housed in an inverse light:dark cycle with constant temperature (68 °F to 72 °F) and humidity (54-59%), and had ad libitum access to water and food.

### 4.2 Video capture, synchronization, and 3D calibration system

Six high-speed monochrome cameras (FLIR CM3-U3-13Y3M-CS 1/2” Chameleon^®^3) were used to record the video data at 100 fps. Based on their location relative to the face, the cameras are labeled LEFT (L), RIGHT (R), TOP LEFT (TL), TOP RIGHT (TR), TOP CENTER (TC) and BOTTOM CENTER (BC) (see **Figure 1b**). The camera location and orientation is selected such that each facial keypoint is in the focused view of at least 2 cameras (see **Supplementary Table 1** for details). The lateral cameras (L, R, TL, TR) are equipped with an 8 mm EFL, *f* /1.4 lens (MVL8M23, Thorlabs) and the center cameras (TC and BC) with a 12 mm EFL, *f* /1.4 lens (MVL12M23, Thorlabs). Lenses are connected to the body of the camera through a C-to-CS-mount (03-618 Edmund Optics) and 3D-printed 1.1 mm (L, R, TL, TR cameras) or brass 2 mm spacer rings (TC, BC cameras) (03-633, Edmund Optics) for fine focal adjustment. The face is illuminated using two infrared lamps (CMVision IR30 WideAngle) with a piece of Kimwipe (Kimtech Science) covering the LED surface acting as a light diffuser to minimize glare.

The six cameras were synchronized using Bonsai (v2.8.1) and an Arduino Mega 2560 REV3, which sends a start signal to Bonsai through the serial port. Upon receiving the trigger signal, Bonsai begins recording frames from all cameras as well as associated metadata for each frame. This hardware signal ensures that different cameras are synchronized at acquisition time. To verify that the camera frames are synchronized post-acquisition, a miniature infrared LED (SML-S13RTT86, Mouser Electronics) is positioned to appear in the field of view of all cameras. As a ground-truth synchronization signal, the LED is on for 10 ms every 10.0 *±* 0.5 sec.

Post-hoc verification of camera synchronization is accomplished by detecting the frame of the rising edge of the LED signal in all views. Using the BC view as a reference, we linearly regress the rising edge times from another view onto the times from the reference view. A perfectly aligned pair of videos should regress onto the identity line, whereas a misaligned view will have a non-zero offset and non-identity slope. The non-zero offset is used as the frame shift to trim the non-reference video, and the slope is used to scale the effective frame rate to match the reference frame rate. This process is repeated systematically for all views and verified using the ground-truth synchronization LED signal.

Video-electrophysiology synchronization is conducted in a similar fashion, with the same synchronization LED signal split and connected to the electrophysiology hardware.

We calibrate camera views using a manufactured calibration board with a standard ChArUco template imprinted on its surface. A vectorized template for the ChArUco board was created using https://github.com/dogod621/OpenCVMarkerPrinter. The template used is for a 7 *×* 7 ChArUco board (4.5 mm marker length, 6 mm square side length, ArUco dictionary DICT_4×4_50). Prior to as well as after recording any experimental data, an experimenter held and rotated the ChArUco board in the focused view of all cameras for at least one minute. These calibration videos were used in Anipose to calibrate the pipeline for triangulation.

### 4.3 Neural network keypoint detections and validations

We utilize video data from across all mice and experimental conditions (feeding experiments, awake recordings from the anesthesia experiment, and recordings from the structure experiment) to train a single DeepLabCut (DLC) model to track 2D keypoints. A total of 491 frames are selected using the K-means clustering algorithm for frame extraction provided by DLC, as well as selected manually (136 frames are manually taken from the feeding experiment). Random uniform sampling is used to separate 20% of the frames for testing, while the remaining 80% are used to train the model. Following the standard guidelines provided by DLC, we select the built-in ResNet-50 model architecture and image augmentation pipeline for our training procedure. The model is trained for 1 030 000 iterations using a learning rate schedule of 0.005 for 10 000 iterations, 0.02 for 420 000 iterations, 0.002 for 300 000 iterations, and 0.001 for 300 000 iterations. After training, the train set error was 2.16 px and the test set error was 4.6 px.

Back-to-back 3D scanner and Cheese3D recordings in anesthetized mice were used to measure the spatial accuracy and resolution of keypoint detection (see **Figure 1**, Supplementary Fig. 1). Each mouse underwent intraperitoneal injection of Ketamine (100 mg kg*^−^*^1^) and Xylazine (10 mg kg*^−^*^1^) cocktail to induce anesthesia, scanned first on the 3D scanner (Einscan-SP, SHINING 3D) and then immediately on the Cheese3D setup. To test the robustness of Cheese3D in detecting 3D keypoints, an alternative set-up was constructed using only four cameras with altered positions and angles (**Supplementary Figure 2d**). The four cameras were equipped with an 8 mm EFL, *f* /1.4 lens (MVL8M23, Thorlabs), a C-to-CS-mount adaptor (03-618 Edmund Optics), and 3D-printed 1.1 mm spacer ring.

### 4.4 Triangulation and 3D tracking optimization

We use the trained DLC model to track keypoints in videos for each camera view separately per experiment. No post-processing is applied to the tracked keypoints. Anipose is used to triangulate 2D keypoints from multiple cameras into a single 3D keypoint per frame. Next, Anipose optimized the 3D keypoint tracking for the full recording by reprojecting the 3D keypoints to 2D in each camera view and minimizing the mean squared error of the reprojected points. Concurrently, the frame to frame velocity of the 3D keypoints is minimized to prevent spurious tracking errors. No post-processing or filtering is applied to the optimized 3D keypoints. To evaluate the performance of the tracking pipeline, we overlaid the optimized 3D keypoints reprojected onto each camera view, and an experimenter curated the accuracy and precision of the tracking results.

### 4.5 Comparison to existing facial motion detection system

The same video data used for Cheese3D was given as input to Facemap. Following Facemap’s guidelines, videos were cropped and some keypoints were discarded or incorporated (i.e.,

Cheese3D whisker pad keypoints were substituted by the base of three whiskers on the right whisker pad, no points on the left whisker pad were labeled, the upper lip keypoints were substituted by a “mouth” middle point and the ears were not labeled). We took the motionless periods (see **Section 4.11** for more information) to quantify 2D jitter. To make the outputs comparable, we reprojected the 3D output of Cheese3D into the six 2D views. We compared the average velocity of the keypoints from each facial region (e.g., eyes, nose) across views per mouse.

### 4.6 Anatomical-based interpretable feature selection

Features are selected and calculated in five tiers with increasing spatial dimension. First, 3D facial keypoints (see **Figure 1**, **Supplementary Figure 1**) are selected based on the following criteria: 1) can be unambiguously and correctly pinpointed by at least three experimenters independently; 2) (for the purpose of 3D calibration) in focused view by at least two cameras; 3) reflect natural facial features and anatomy. Second, Euclidean distances between 3D keypoints within a localized facial region (e.g. the left eye) are calculated; Third, areas are calculated for the sets of keypoints that form a closed polygon; these include the eye, ear, and mouth areas. Fourth, the angle between the ear and snout is calculated as a measure for how forward-orienting the ears are with respect to the whole face. Fifth, the volumes of the nose bulge and whisker pad bulge are calculated to reflect anatomically relevant volumes [5].

The area of the eye and ear groups are calculated based on a flattened 2D ellipse. Each group consists of four points defining the major and minor axis endpoints of the ellipse. Since all four points are not necessarily coplanar, we assume that the ellipse can be bent along the minor axis. To compute the area of this bent ellipse, we begin by defining the major axis (using the front and back of the eye or the base and tip of the ear). Next we compute the midpoint of the major axis and calculate the Euclidean distance from this midpoint to each of the remaining two minor axis endpoints. The sum of these two distances defines the length of the minor axis after a potential bend has been flattened. Using the major and minor axis lengths, we compute the final ellipse area as the standard area of a 2D ellipse in Euclidean space. The area of the mouth can be computed as the standard area of a triangle in Euclidean space. The right and left upper lip points and one central lower lip point form the vertices of the triangle. The volume of the nose bulge is calculated for an irregular tetrahedron defined by the nose top, left and right pad top, and the midpoint between the front of the eyes. We use the standard volume for an irregular tetrahedron in Euclidean space. The volume of the whisker pad bulge is calculated for an irregular pyramid defined by the nose bottom, left and right pad top, and left and right pad side points. We compute the convex hull defined by these points, then calculate the volume of the hull by dividing the hull into smaller tetrahedrons. The specific choice of tetrahedrons used is determined by the SciPy library.

### 4.7 Analysis of keypoint jitter

We quantified the tracking jitter of 3D keypoints and facial features using a five-minute video segment where the experimenter identified no discernible movement (referred to as the ’motionless period’). Next, we calculated the magnitude of the frame to frame velocity of each keypoint during the selected periods which we refer to as the jitter velocity of a keypoint. We use frame to frame velocity as our metric for jitter so that we focus on short time scale noise in the tracking instead of slow moving trends in the tracking that may occur over minutes or hours. To visualize the distribution of keypoint jitter velocity in **Figure 2a**, we compute a Gaussian kernel density estimate (KDE) using the histplot function in the Seaborn plotting library (v0.13.2). The bin size is set to 0.05 mm/sec, and the KDE bandwidth is set using the scotts_factor function in the SciPy library (v1.10.1). We summarize the distribution of jitter velocity during the motionless period by computing the average velocity over the entire period per mouse in **Figure 2b**.

To assess how the jitter velocity of keypoints affects our anatomical features, we computed the absolute frame to frame velocity of each feature during the selected periods which we refer to as the jitter velocity of an anatomical feature. We selected the 99.9th-percentile of the anatomical jitter velocity distribution per mouse as our motion threshold. Any movement with a frame to frame velocity below this threshold will be considered noise. The motion threshold across mice is summarized in **Supplementary Figure 3**.

### 4.8 Headpost design and surgery

The custom-designed stainless steel headpost for head-fixation consists of a 6 mm *×* 4 mm *×* 1 mm rectangular base and a small 10 mm *×* 3 mm post that fits into the headpost holder. A groove was added on each lateral end of the base design to facilitate metabond adhesion during implant surgery. The headpost has a conical notch etched on the side to secure in the headpost holder with a screw fastener. The headpost holder is angled at 27.9*^◦^* following observation of the natural head angle of mouse eating to maximize comfort.

To implant the headpost, 2-month-old mice were anesthetized with isoflurane (SomnoFlo, Kent Scientific; 3–5% induction, 1–2% maintenance). Once anesthetic depth was achieved, mice were placed onto a stereotaxic apparatus where body temperature was maintained using a heating pad. After flattening the skull using skull landmarks, the base of the headpost is positioned above the medial-lateral midline, and immediately anterior to lambda, and secured using adhesive cement (Metabond, C&B). Following surgery, animals were administered buprenorphine (0.1 mg kg*^−^*^1^) and allowed to recover on a heating pad before returning to their home cages, where the mice continue to recover for one week before being acclimated to sitting in a tunnel and head-fixation for one to two weeks.

### 4.9 Electroencephalography surgery and data acquisition

One day before surgery, a biotelemetry unit (HD-X02, Data Sciences International) was thoroughly disinfected in CIDEX OPA solution (Advanced Sterilization Products). Surgery procedure is as described above, with 2 stainless steel screws (00-96 *×* 1/16, EEG, IROX screw, Data Sciences International) inserted through craniotomy as cortical electrodes (Screw 1:1.5 mm anterior of bregma, 1.5 mm lateral of midline to the left; Screw 2: 1.5 mm posterior of bregma, 3.5 mm lateral of midline, contralateral to screw 1. An incision was made to expose the upper trapezius muscles on the animal’s back to place the transmitter into the subcutaneous pocket. A pair of EEG leads was attached to the cortical screws and secured with a small amount of dental cement, and a pair of EMG leads was threaded through the upper trapezius muscles (one on each side of the midline) and held in place by 6-0 polyamide sutures. Tissue adhesive (3M Vetbond) was applied to the skull, before attaching the headpost (described above). Post-surgery procedures were same as descibed above, with at least two weeks recovery time before experiments.

Recording was conducted using a wireless recording system (Data Sciences International): Transmitters communicated with PhysioTel receivers connected to PhysioTel Matrix MX2 acquisition interface. An Arduino MEGA generated a square signal that was synchronously sent to an infrared LED, visible in the Cheese3D camera system, and the PhysioTel Signal Interface, connected to the MX2 acquisition interface. The acquisition interface communicates with the EEG recording computer running Ponemah 6.5. All hardware devices are configured within Ponemah to record EEG, EMG, and LED synchronizing signal at 1 kHz, and body temperature at 1 Hz. After the experiment, data was exported to CSV format using NeuroScore 3.3.1 - Build 9317 (Data Sciences International).

### 4.10 In vivo electrophysiology recording and electrical stimulation surgery and data acquisition

Surgery was conducted as described above. A craniotomy was performed (anterior to bregma 1.5 mm, lateral 1.0 mm) to insert a grounding pin (male connector pin, A-M systems, Sequim, WA, USA) at a 45*^◦^* angle, with the tip of the pin pointing at the rostral direction. The skull was sealed with a tissue adhesive (3M Vetbond, St. Paul, MN, USA) before the head gear implant was attached and further secured with dental cement (C&B metabond, Parkell, Brentwood, NY, USA). The rest of the exposed skull was covered with additional dental cement to further secure the head gear in place. Post-surgery procedures were same as descibed above, with at least one week recovery time before experiments water deprivation to aid acclimation (one to three weeks) to awake head-fixation.

Following acclimation, a 1 mm diameter brainstem cranial window was made above (posterior to lambda 1.8 mm, lateral 1.25 mm). Head-fixed mice were then stimulated under ketamine and xylazine anesthesia for 3-5 sessions (“stimulation sessions”), followed by 2 awake acute recording sessions (“recording sessions”). The camera configuration differs slightly from other datasets—the lateral and top center cameras (L, R, TL, TR, TC) are equipped with an 8 mm EFL, *f* /1.4 lens (MVL8M23, Thorlabs) and the bottom center camera (BC) with a 12 mm EFL, *f* /1.4 lens (MVL12M23, Thorlabs). In the first stimulation session, multiple brainstem locations were probed with a stimulation grid search. In subsequent stimulation sessions, a 32-channel four-shank silicon probe (A4*×*8-5 mm-100-200-177, NeuroNexus Technologies) was inserted at the mapped locations from the first session. Single electrical pulses (2 ms pulse duration, biphasic) were delivered to either the entire probe or specific divisions of individual shanks via Allego software (NeuroNexus Technologies, USA) every 2 sec to 5 sec. In recording sessions, a 32-channel single-shank silicon probe with 46 kΩ to 54 kΩ impedance (H7b or H8b, Cambridge NeuroTech) was inserted into the brain region mapped during the previous stim sessions. The probe was coated with lipophilic dyes DiI or DiO (10% w/v) to reveal the probe track post-hoc. Recordings began at least 15 min after probe insertion to ensure recording stability. Voltage signals were amplified using an RHD2132 amplifier (Intan Technologies) and acquired at 30 kHz with a NeuroNexus XDAQ ONE system. After the recording, single electrical pulses (0.5 µA to 2 µA) were delivered to all sites on the probe to induce facial movements to verify probe placement location.

Initial spike sorting was performed using Kilosort 2.5 with default parameters, followed by manual curation in Phy. Clusters with inter-spike interval violations, low signal-to-noise ratios, or low stability through the recording session were excluded from single-unit analyses.

### 4.11 Analysis of kinematics during anesthesia

For the anesthesia experiments (see **Figure 2**), awake spontaneous movements were recorded in Cheese3D for 5 min, followed by intraperitoneal injection of Ketamine (100 mg kg*^−^*^1^) and Xy-lazine (10 mg kg*^−^*^1^) cocktail to induce anesthesia, before returning to Cheese3D to record facial movement during and recovery from anesthesia. Temperature was maintained on a heating pad, and the exact time of injection was recorded.

To measure the wakefulness of each mouse during anesthesia, we compute the magnitude of the frame to frame velocity of each anatomical feature over the entire recording. We labeled each time point as movement if the frame to frame velocity crosses the previously computed motion threshold, while time points where the velocity is below the threshold is labeled as no movement. **Figure 3a** shows an example raster plot of time points labeled as movement for one mouse.

We analyzed slow drift of the anatomical and EEG features during anesthesia using a moving average of each feature during the entire recording period. The moving average is computed using a 10 sec (facial features) or 40 sec (EEG features) wide sliding window average. **Figure 3d** shows exemplar filtered features for one mouse over the entire recording period. We visualized the filtered features across all mice and selected three representative features across three facial regions—left ear angle, left eye height, and nose bulge. We trained a model across mice to predict time since injection using the selected features during anesthesia. Our model’s input consists of quadratic terms of the feature at the current time point and initial time point (quadratic terms computed using Scikit Learn’s (v1.4.2) PolynomialFeatures class) as well as a constant bias. We performed a linear regression from our quadratic input terms to the current time since injection using the Lasso class from Scikit Learn (v1.4.2) where we used leave-one-out cross-validation and grid search (over 100 values from 0.1 to 100) to find the optimal regularization coefficient. A separate model was trained for features from the facial regions and the EEG power bands. From a total of 12 sessions (across *n* = 3 mice), we held out 3 random sessions for testing and used 9 for training, repeating this procedure 220 times. We assessed the performance of each model by predicting the time since injection for each test session. A moving average filtered (using the same filter as **Figures 3c**, **3d**) prediction for a single mouse and exemplar feature sets is shown in **Figure 3e**. We computed the average root mean squared error of the three sessions for each of the 220 runs in **Figure 3f**.

### 4.12 Analysis of chewing kinematics

FED3 [45] was used to dispense chocolate-flavored 20 mg pellets (Dustless Precision Pellets, F05301, Bio-Serv) on demand during the feeding experiment (see **Figure 4**). A funnel and tubings are placed underneath the FED3 spout to collect the dispensed pellet and deposit it on a translucent plastic spoon (Measuring Scoop S378, Parkell). The spoon was attached to a servo motor connected to a 3D printed linear actuator to bring the pellet to the mouth, and then retracted to await the next pellet. Animals in the feeding experiments were gently food-restricted and acclimated for two days to eating from the spoon while head-fixed, to facilitate food consumption during the experiment. Each mouse was recorded eating 10 to 13 pellets in one session, and allocated 30 sec per pellet. Dropped pellets were excluded from subsequent analysis.

We distinguished the ingestion and mastication phases of chewing based on the shape of the lower envelope of the mouth area during the consumption of each pellet per mouse. An example lower envelope is shown in **Figure 4c**. To compute the envelope, we invert the mouth area by negating it, then identifying the peaks of the negated signal using the find_peaks function in SciPy (v1.10.1) with a window of 200 m sec. The lower envelope is defined by linearly interpolating the calculated peaks, then median filtering the interpolated curve with a window of 1.49 sec. We defined the transition time from ingestion to mastication as when the lower envelope drops sharply as shown in **Figure 4c**. To quantify the time when the envelope drops, we computed the cumulative area under the envelope during the consumption of each pellet. The cumulative area quickly increases during ingestion, then sharply transitions to a slower increase during mastication. The “knee” in the cumulative area under the envelope was used to quantitatively define the transition time. We used the Kneedle algorithm (with the sensitivity parameter set to 1) to identify the knee point (transition time) for each pellet per mouse shown in **Figure 4d**. The Python kneed (v0.8.5) library was used as our Kneedle implementation.

In **Figures 4e**, **4f**, we compared the mouth area and nose bulge during the consumption of pellets by z-scoring each anatomical feature separately per pellet per mouse. For **Figure 4f**, we plot the normalized mouth area and nose bulge against each other for an example mouse where each point constitutes a single frame. We color each point based on whether it occurs before or after the transition time for the corresponding pellet.

We defined the eye protrusion in **Figures 4h**–**4k** as the Z coordinate of the left eye back keypoint (we observed similar behavior for the right eye back). To quantify the degree of co-ordination between the mouth area and eye protrusion, we z-scored each feature per pellet per mouse. Next, we computed the cross-correlation between the normalized features per pellet per mouse separately for the ingestion and mastication phases. **Figure 4i** shows the mean cross-correlation taken across pellets for a single mouse. We identified the peak cross-correlation by selecting the time point with the largest absolute cross-correlation per pellet per mouse as shown in **Figures 4j**, **4k**.

### 4.13 Analysis of in vivo electrical stimulation and electrophysiological recording

Following data collection as described in **Section 4.10**, video and ephys data was synchronized as described in **Section 4.2**. To obtain stimulus triggered facial responses in **Figures 5b**–**5f**, a stimulus trigger signal aligning with the pulse duration was recorded on an analog channel of the electrophysiology data and used to drive an LED visible in the video data. For all but one sessions, the rising edge of the recorded trigger signal in the electrophysiological data was used as the timepoint for stimulus triggered alignment. In one session, the analog channel was not available due to hardware failures, and the LED in the video was used instead.

Next, we computed the peak displacement of various facial features from their pre-stimulation baseline value. For 100 ms before the stimulation trigger, we measure the average value of a facial feature, then we define the peak displacement as the maximum (positive or negative) deviation away from the baseline average during a 100 ms period after the stimuluation trigger. This is done on a trial by trial basis for each stimulation. In **Figure 5b** (top right), we plot the absolute value of the peak displacement for the eye height, taking the mean and standard deviation across trials of the same stimulation current amplitude.

We also evaluated our ability to distinguish stimulation-triggered movement from noise using receiver operating characteristic (ROC) curve in **Figure 5b** (bottom right). For a given stimulation current, taking a 100 ms before stimulation as a “false positive” response (noise) period and 100 ms after stimulation as a “true positive” response (stimulation-triggered movement) period, we measure detected responses whenever the minimum ipsilateral eye height during the period is below a threshold. Sweeping the threshold from 10% below the minimum to 10% above the maximum eye height, we quantify the response rate during each period as the false positive rate and true positive rate for every threshold. From this, we construct an ROC curve, and measure the area under the curve using NumPy’s trapz function. The ROC-AUC for each amplitude is shown in **Figure 5b**.

In **Figure 5d**, we show the absolute peak displacement of several facial features at 10 µA across all stimulation locations. We omitted some locations (appearing as white blank squares in the figure) when the peak displacement amplitude was below the jitter threshold for that feature as defined in **Section 4.7** and shown in **Supplementary Figure 4**.

For in vivo electrophysiological recordings, spike trains were shuffled 1000 times using a cyclic shuffling method, in which the entire spike train was shifted by a time within *−*60 sec to *−*15 sec or 15 sec to 60 sec [46], with shifted spikes outside the time boundary wrapped to the start or the end of the time duration. To examine the correlation between neural activity and facial movements, actual or shuffled neural firing rates binned at 10 ms were cross-correlated with each facial feature. If the peak of the cross-correlation lies outside the 99.9% confidence interval of the shuffled cross-correlation, and the corresponding lag is within *−*100 ms to 0 ms, the unit is considered statistically correlated with the facial feature (p-value *<* 0.001 as set by 1000 shuffles). The significance of facial feature tuning was further assessed using 95% confidence intervals of shuffled tuning curves and by manually verifying tuning specificity through facial video recordings overlaid with spike sounds.

### 4.14 Data and Code Availability

Code to reproduce the reported results is available at https://github.com/Hou-Lab-CSHL. Data presented in this paper are available upon requests from the corresponding author. Following acceptance of the manuscript they will be archived in a permanent public repository.

## Acknowledgements

We would like to thank Y. Zhang for double-blinded 3D scan annotation, R. Eifert for manufacturing ChArUco board and headpost, X. Zhang for assembling FED3, S. Rahman for double-blinded measurements, J. Merrill and S. Lyons of Animal Imaging Core for microCT imaging. L. Corcos for video editing, CSHL facilities for constructing the behavioral experiment room. In addition, we thank M. Bagnall, L. Boyd, B. Cowley, D. Klindt, B. Sabatini, and S. Shea for comments on the manuscript. This work was funded by a Brain and Behavior Research Foundation NARSAD grant to X.H.H., a CSHL NeuroAI Scholarship funded by the Schmidt Foundation to K.D., and a Fulbright Scholarship to I.N.M.

## Supplementary Information

### Supplemental Figures

**Supplementary Figure 1:**
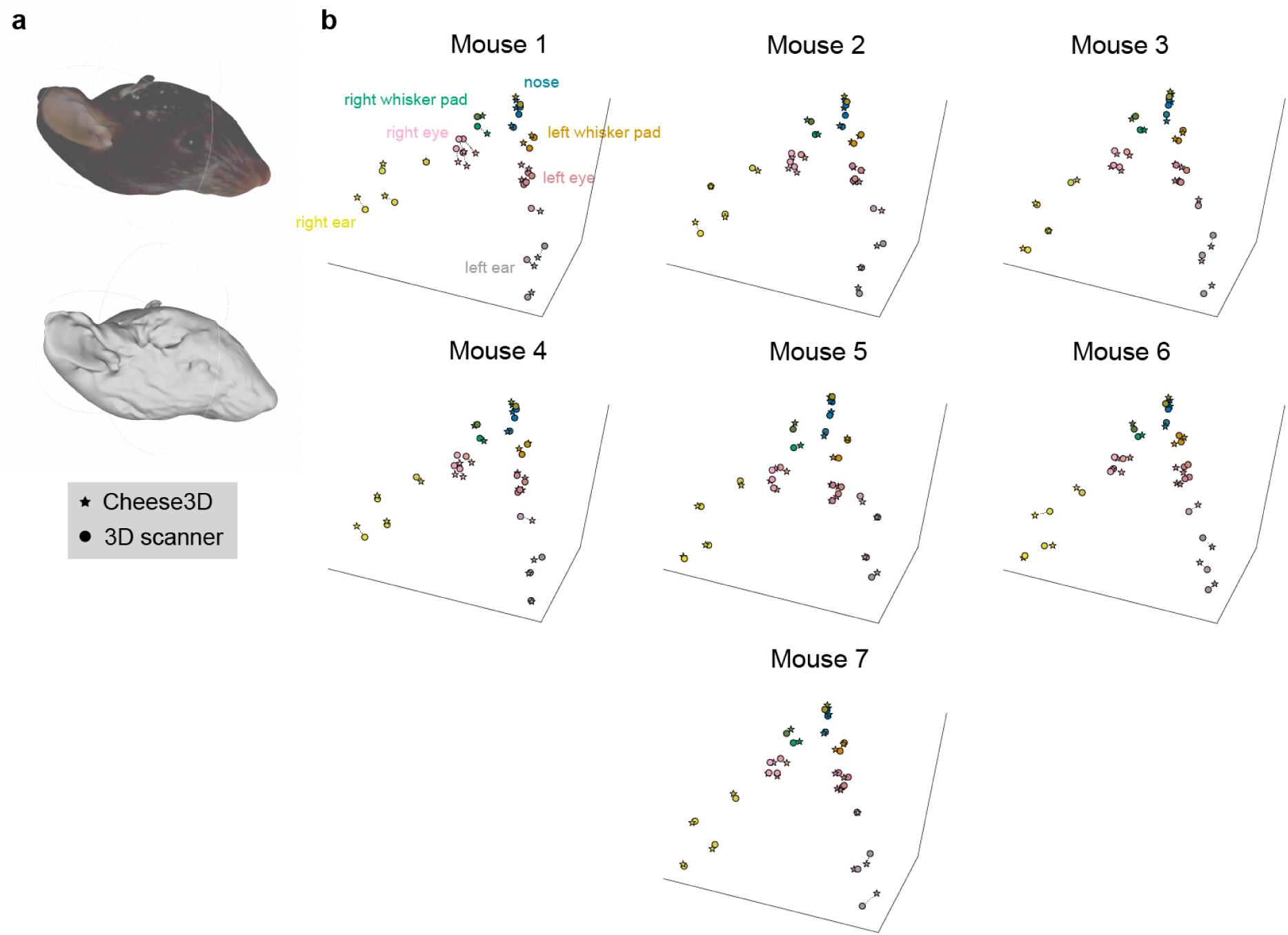
3D facial keypoint comparison between Cheese3D and 3D scanner **(a)** Example textured (above) and untextured (below) mesh of mouse face obtained in 3D scanner used for validating Cheese3D keypoint placement. **(b)** 3D scatter plot showing corresponding 3D scanner (solid circle) and Cheese3D (star) triangulated keypoints for all mice shown in Figure 1i.

**Supplementary Figure 2:**
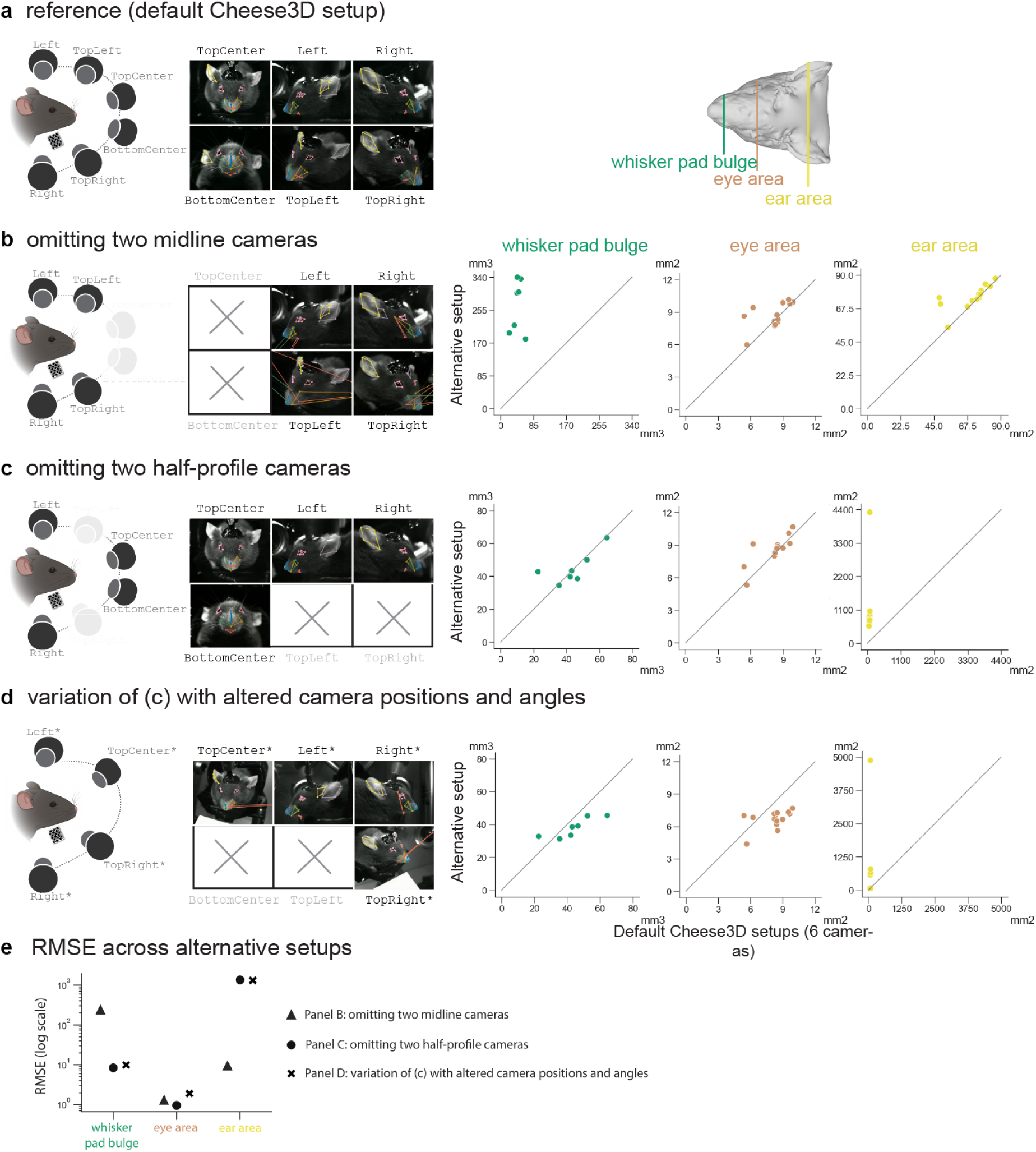
Utility and necessity of six cameras in capturing mouse face **(a)** Default setup for Cheese3D with a calibrated array of six (three pairs of) cameras: hardware schematic (*Left*) and 3D facial keypoints from the Cheese3D model projected onto example video frames (*Right*), same as Figure 1. **(b)** Omitting a pair of midline cameras results in distorted keypoint inference and measurements of midline structures (Whisker pad bulge RMSE: 231.92 µm^3^) but intact lateralized structures (Eye area RMSE: 1.27 µm^2^; Ear area RMSE: 9.22 µm^2^) compared to the default Cheese3D setup (Figure 1i). **(c)** Omitting a pair of half-profile cameras results in altered keypoint inference and measurements of the most lateralized structures (Ear area RMSE: 1349.22 µm^2^) but intact midline structures (Whisker pad bulge RMSE: 8.34 µm^3^) and eye area (RMSE: 0.95 µm^2^) compared to the default Cheese3D setup. **(d)** A variation of setup of **(c)** with altered camera positions and angles does not rescue keypoint inference and measurements of the most lateralized structures (Ear area RMSE: 1311.45 µm^2^) but has similarly intact midline structures (Whisker pad bulge RMSE: 9.89 µm^3^) and eye area (RMSE: 1.90 µm^2^). **(e)** Summary of RMSE for various facial regions across **(b)**, **(c)**, and **(d)**.

**Supplementary Figure 3:**
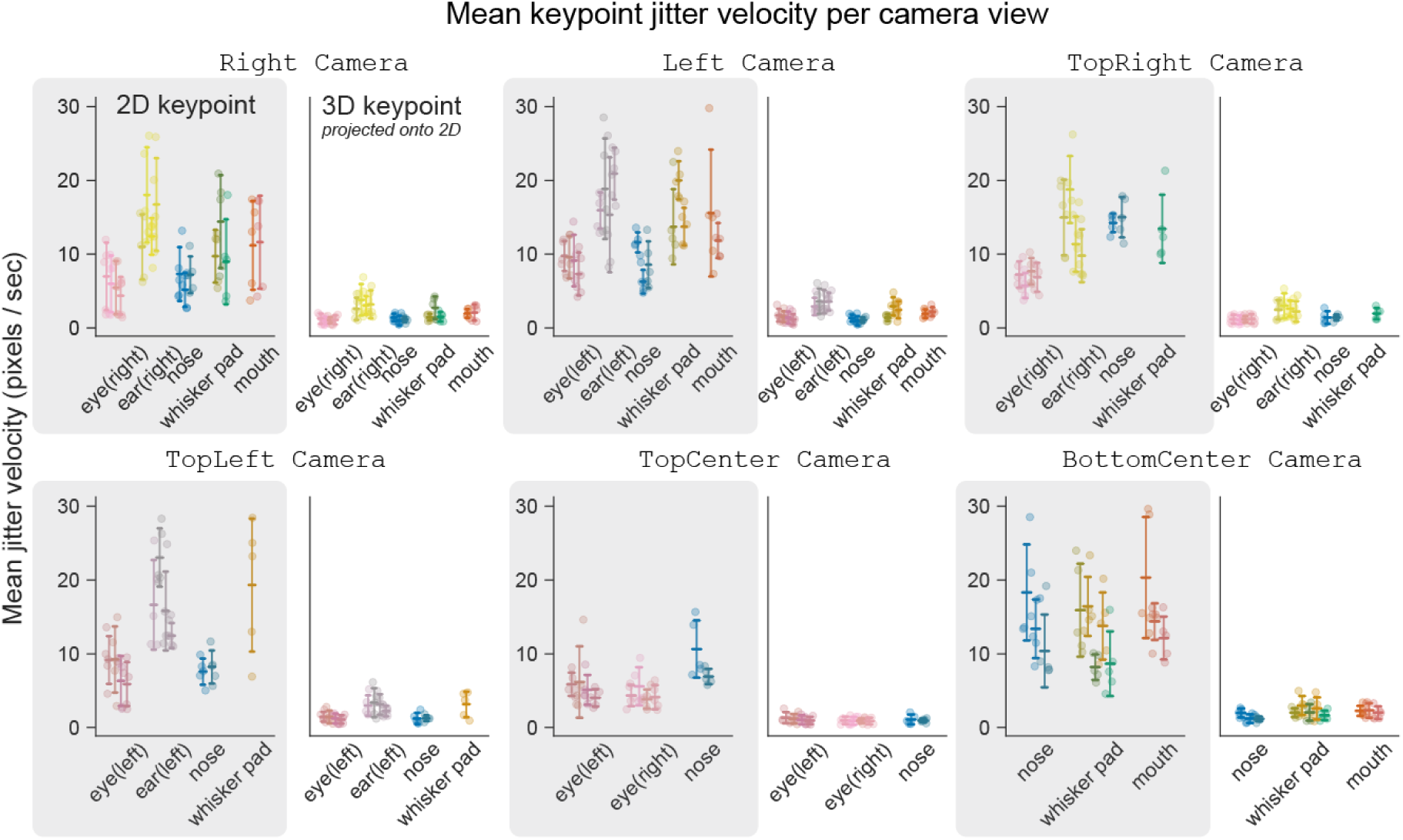
Keypoint jitter comparison between 2D and 3D Summary of keypoint-specific jitter where each pair of panels represents a different camera view, and each pair shows the jitter pre- (shaded) and post-triangulation (Mean jitter velocity, pre-triangulation. Ear (left): 17.37 *±* 5.86 px/sec, Ear (right): 14.12 *±* 5.26 px/sec, Eye (left): 7.29 *±* 3.39 px/sec, Eye (right): 5.67 *±* 2.67 px/sec, Nose: 10.05 *±* 4.68 px/sec, Whisker pad: 13.55 *±* 5.86 px/sec, Mouth: 13.86 *±* 6.12 px/sec; Mean jitter velocity, post-triangulation. Ear (left): 3.16 *±* 1.37 px/sec, Ear (right): 2.85 *±* 1.50 px/sec, Eye (left): 1.29 *±* 0.67 px/sec, Eye (right): 1.02 *±* 0.50 px/sec, Nose: 1.23 *±* 0.55 px/sec, Whisker pad: 2.21 *±* 1.15 px/sec, Mouth: 2.10 *±* 0.74 px/sec; all *p <* 0.0001, one-sided Wilcoxon matched-pairs test (pre-triangulation *>* post-triangulation)).

**Supplementary Figure 4:**
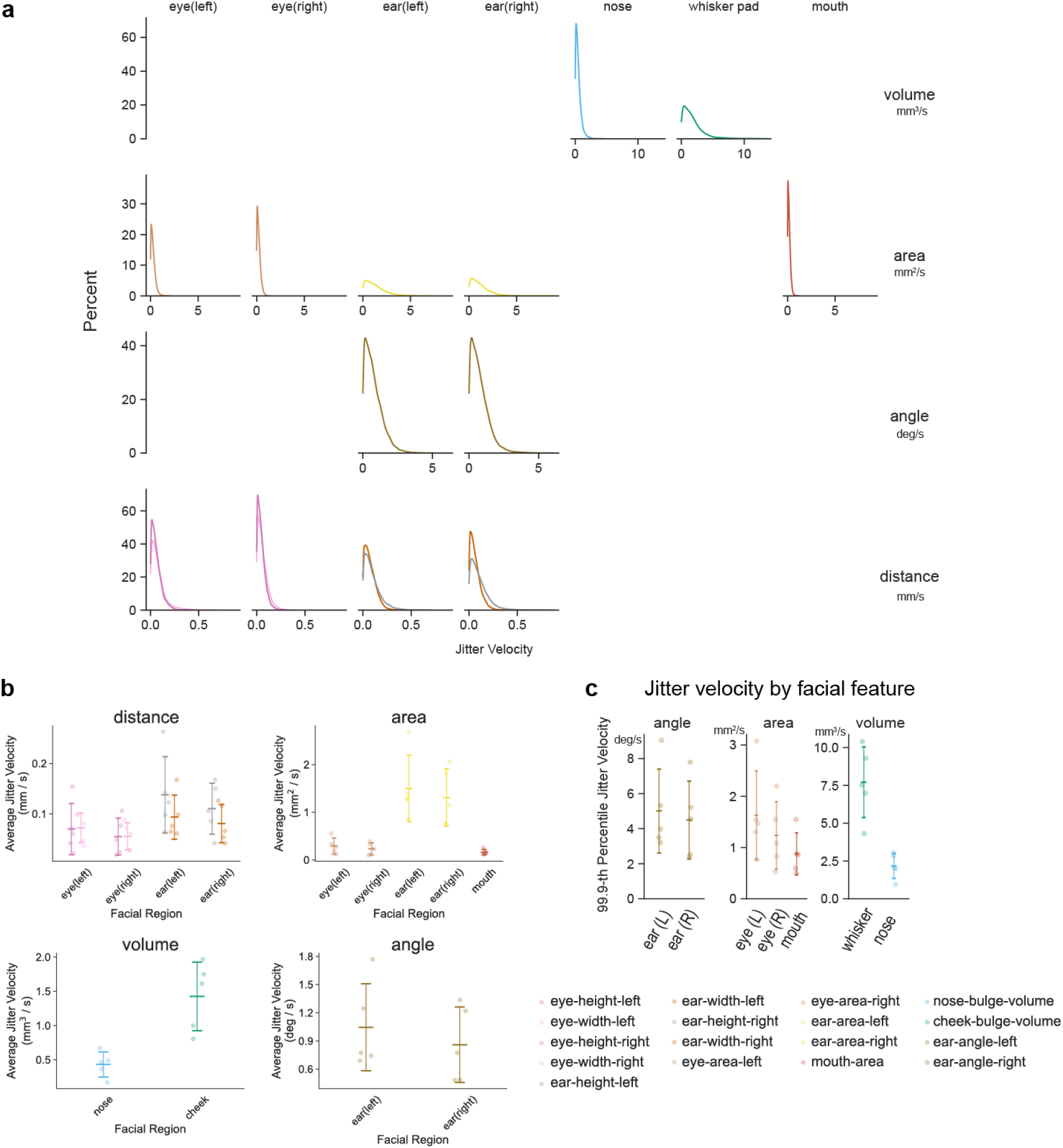
Tracking jitter by 3D facial feature **(a)** Distribution of facial feature-specific jitter (frame-to-frame velocity) relative to the mean jitter during a motionless period for an example mouse. Each column indicates a different facial region, and each row indicates a different type of measurement. **(b)** Summary of facial feature-specific jitter where each panel represents a different type of measurement. **(c)** Summary of 99.9-th percentile jitter velocity used in Figures 3a, 3b.

**Supplementary Figure 5:**
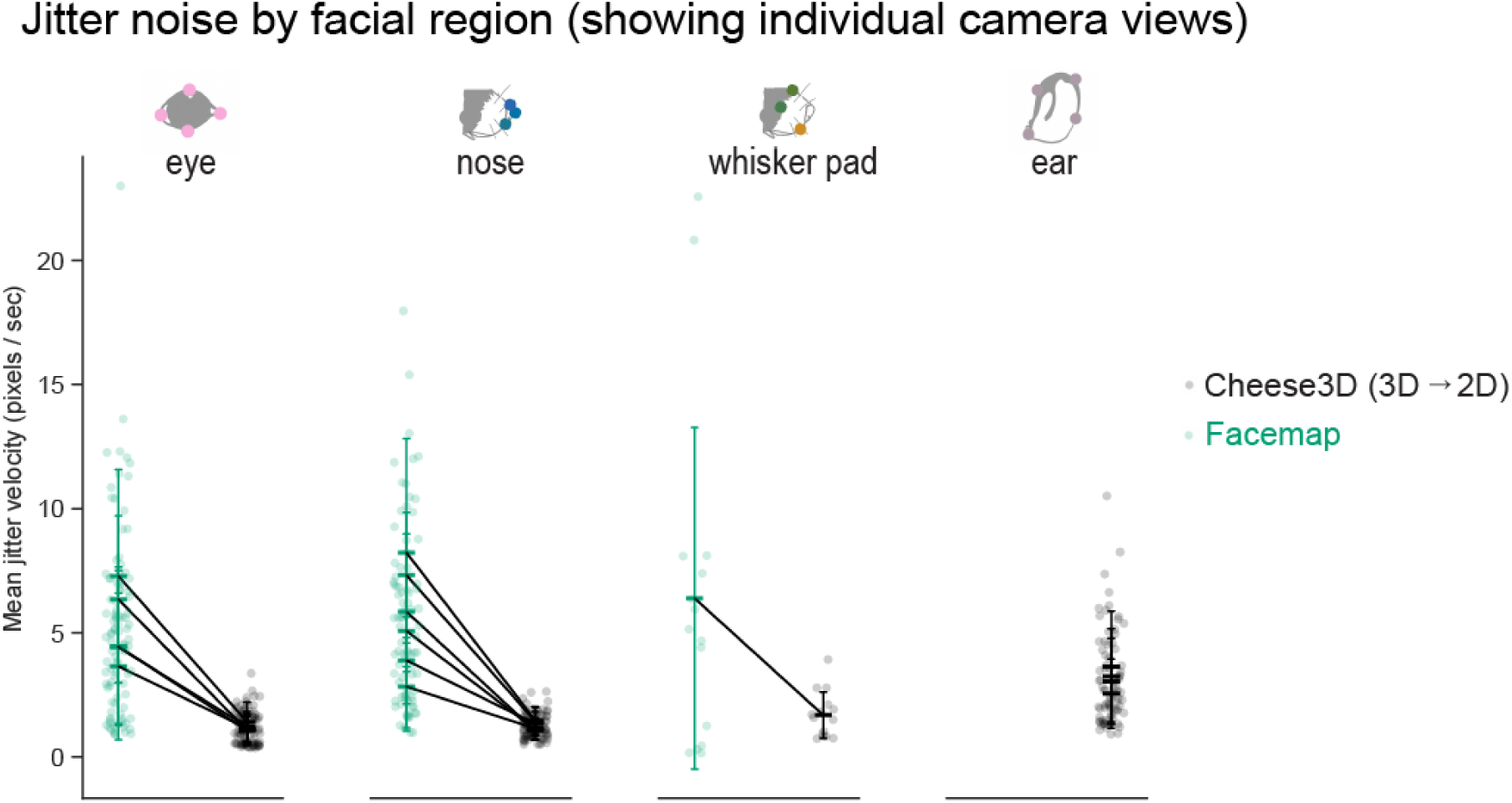
Reduction in keypoint tracking jitter from Facemap to Cheese3D by facial region Comparison of jitter between Facemap’s output and Cheese3D’s output. Each datapoint represents the mean jitter velocity of a single mouse for a particular facial region and the camera views where the region is visible from. (Mean *±* std jitter velocity, Facemap. Eye (Left view): 7.29 *±* 4.29 px/sec, Eye (Right view): 4.46 *±* 3.19 px/sec, Eye (TopRight view): 3.65 *±* 2.95 px/sec, Eye (TopCenter): 6.35 *±* 3.36 px/sec, Eye (TopLeft): 4.42 *±* 3.09 px/sec, Nose (Left view): 5.08 *±* 3.90 px/sec, Nose (Right view): 2.83 *±* 1.77 px/sec, Nose (TopCenter view): 5.85 *±* 2.41 px/sec, Nose (TopRight view): 8.23 *±* 4.59 px/sec, Nose (TopLeft view): 3.89 *±* 1.75 px/sec, Nose (BottomCenter view): 7.32 *±* 2.51 px/sec, Whisker Pad (Right view) 6.39 *±* 6.87 px/sec; Mean jitter velocity, Cheese3D. Eye (Left view): 1.41 *±* 0.80 px/sec, Eye (Right view): 1.07 *±* 0.57 px/sec, Eye (TopRight view): 1.15 *±* 0.60 px/sec, Eye (TopCenter view): 1.07 *±* 0.59 px/sec, Eye (TopLeft view): 1.22 *±* 0.61 px/sec, Nose (Left view): 1.10 *±* 0.42 px/sec, Nose (Right view): 1.15 *±* 0.42 px/sec, Nose (TopCenter view): 1.05 *±* 0.34 px/sec, Nose (TopRight view): 1.48 *±* 0.55 px/sec, Nose (TopLeft view): 1.34 *±* 0.48 px/sec, Nose (BottomCenter view): 1.43 *±* 0.58 px/sec, Whisker Pad (Right view): 1.69 *±* 0.92 px/sec, Ear (Left view): 3.64 *±* 2.24 px/sec, Ear (Right view): 3.25 *±* 1.91 px/sec, Ear (TopRight view): 2.55 *±* 1.39 px/sec, Ear (TopLeft view): 3.04 *±* 1.74 px/sec; all *p <* 0.005, one-sided Wilcoxon matched-pairs test (pre-triangulation *>* post-triangulation)).

**Supplementary Figure 6:**
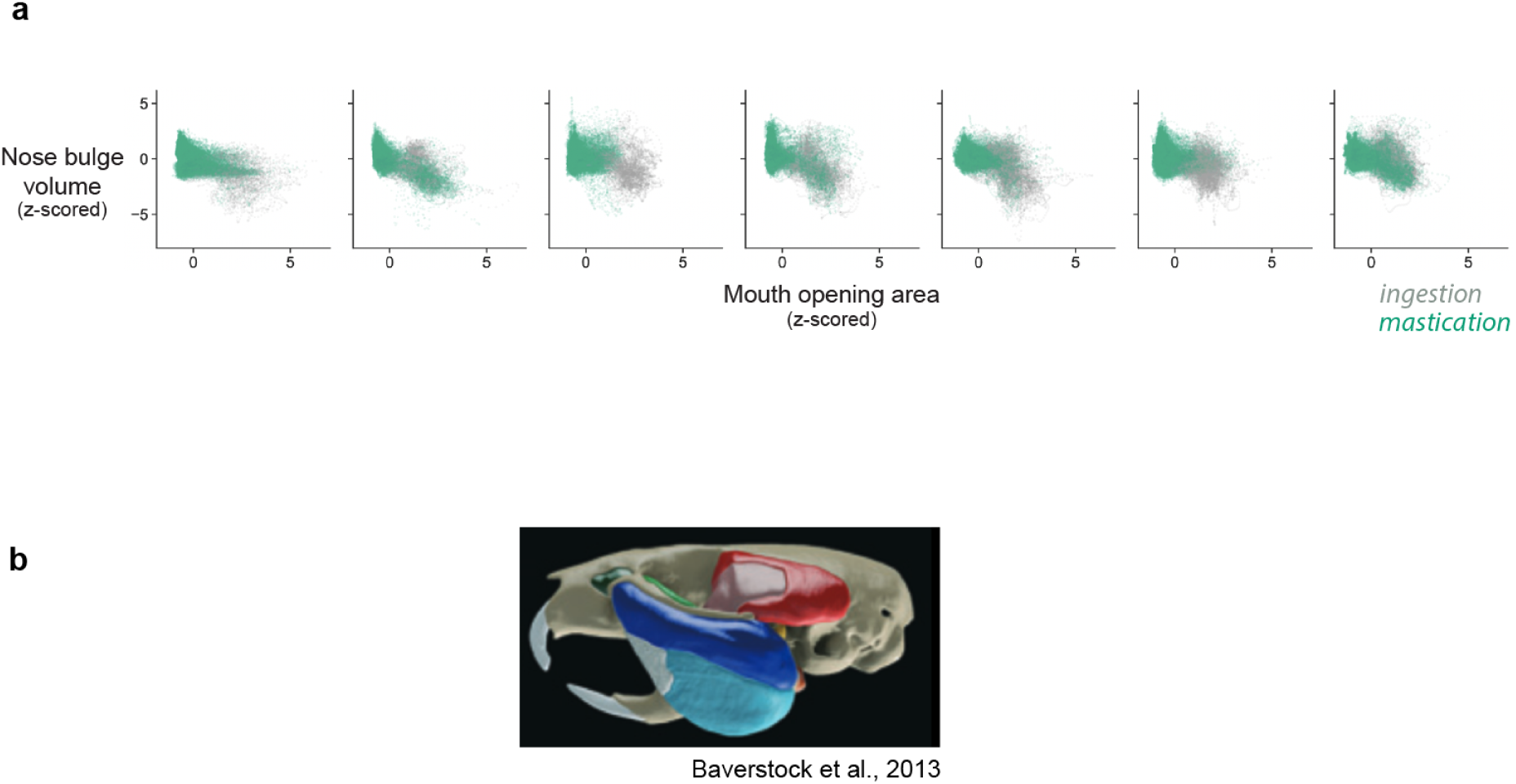
Visualizing putative division between ingestion and mastication across mice. (**a**) Mouth opening area and nose bulge volume (z-scored) scatter plot across mice, sorted by ascending putative transition time between ingestion (gray) and mastication (green) (see Figure 4d). The example mouse shown in Figure 4f is second from the right. **(b)** Segmentation of muscles of mastication shown wrapping around the eye socket [26].

**Supplementary Figure 7:**
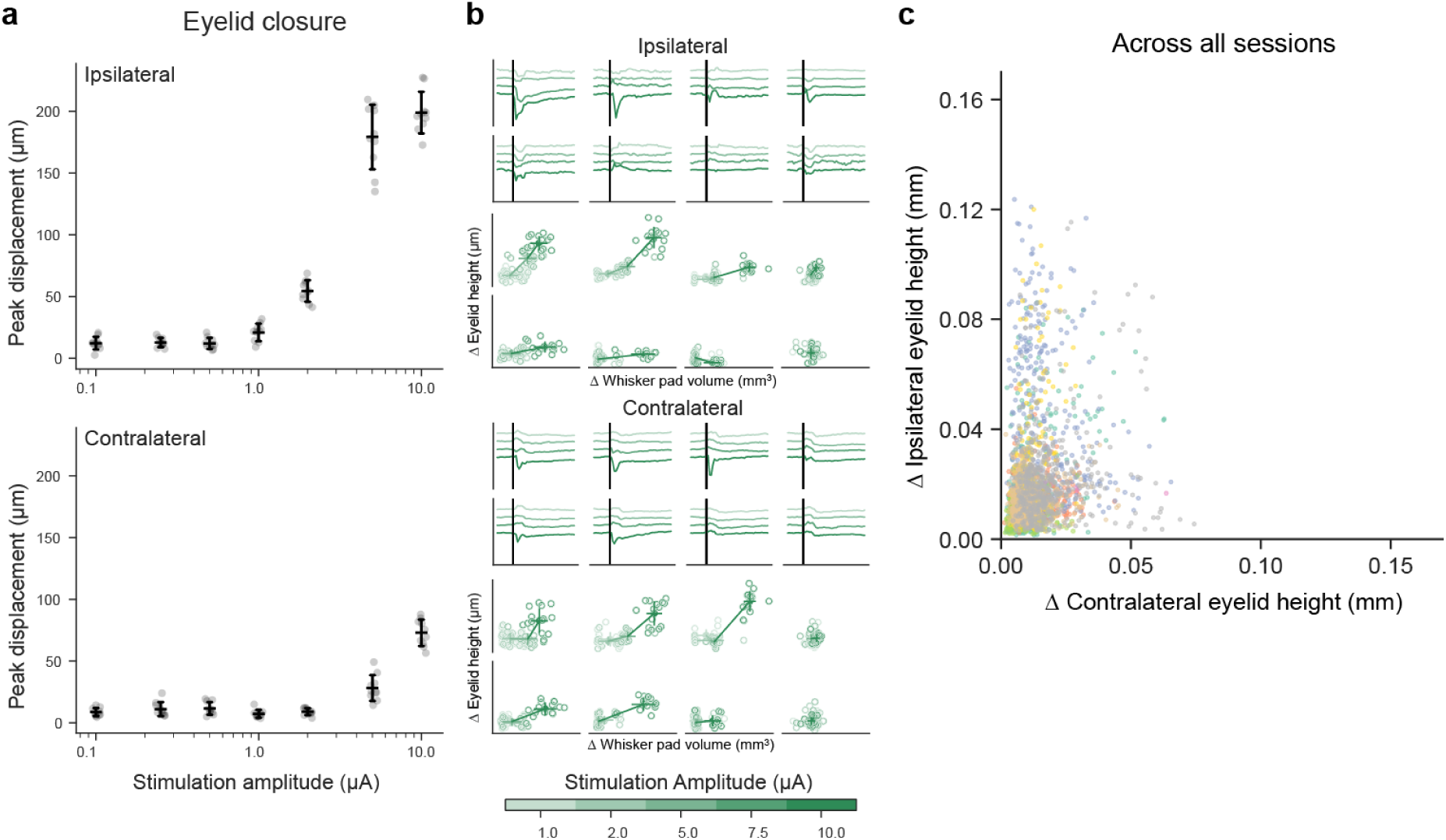
Ipsilateral vs. contralateral asymmetry of eletrically stimulated facial movement. (**a**) Peak absolute deflection of ipsilateral eyelid closure (top) and contralateral eyelid closure (bottom) measured as a function of stimulation current amplitude. **(b)** Top half, from top to bottom: Ipsilateral eyelid closure responses to spacial-specific focal stimulation at varying amplitude; each trace indicates the mean response per amplitude (*n* = 15-30 stimulations per amplitude and location). Pairwise responses of ipsilateral eyelid closure vs. whisker pad volume responses. Bottom half: same as top but for contralateral eyelid responses. **(c)** Pairwise relationship of peak absolute deflections of responses for ipsilateral vs. contralateral eyelids, but colored by experimental sessions (9 sessions from *n* = 4 mice).

### Supplemental Videos

**Supplementary Video 1:**
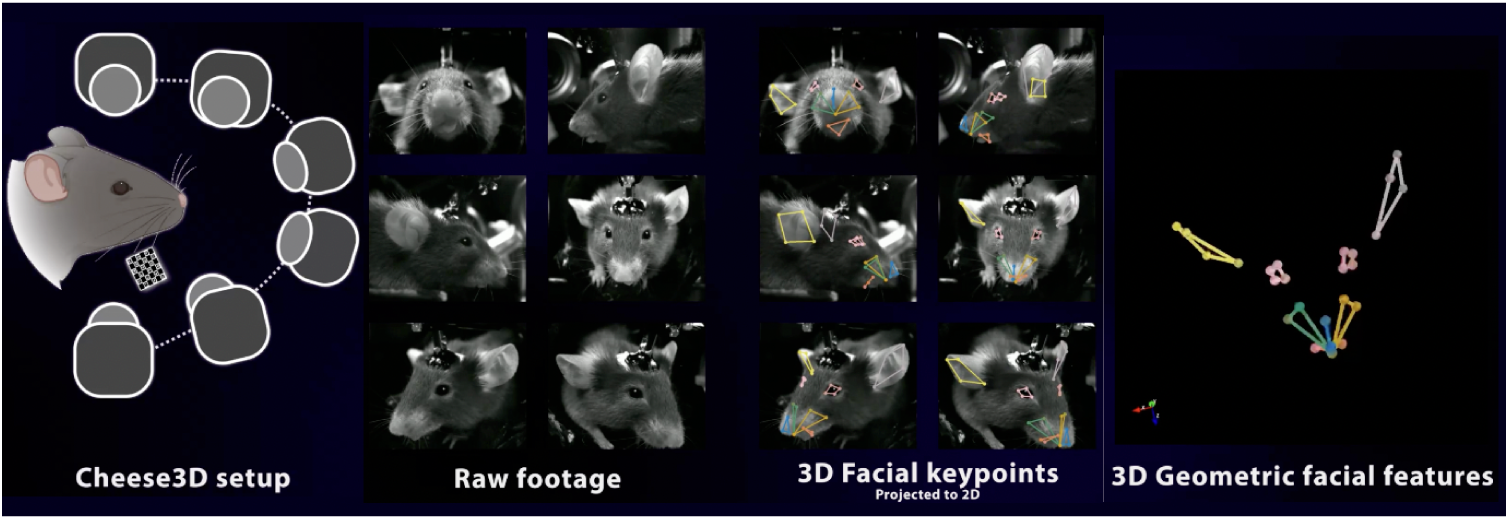
Cheese3D tracks whole-face movement in mouse

**Supplementary Video 2:**
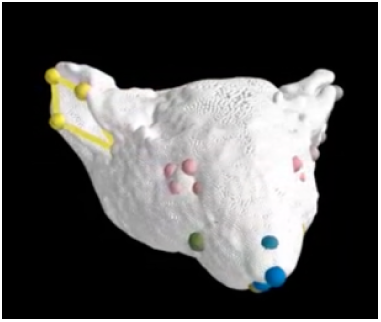
Facial keypoints from Cheese3D model overlaid on mesh from 3D scanner

**Supplementary Video 3:**
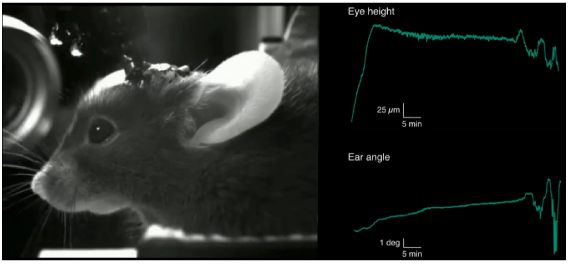
Example of gradual change in eye height and ear angle during anesthesia (sped up 500*×*)

**Supplementary Video 4:**
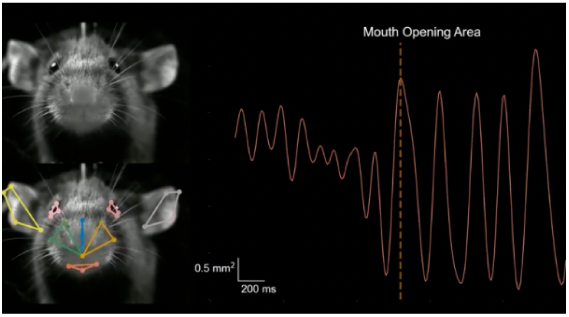
Example transition from ingestion (using incisors) to mastication (using molars) as capitulated in mouth opening area in 3D (slowed down 4*×*)

**Supplementary Video 5:**
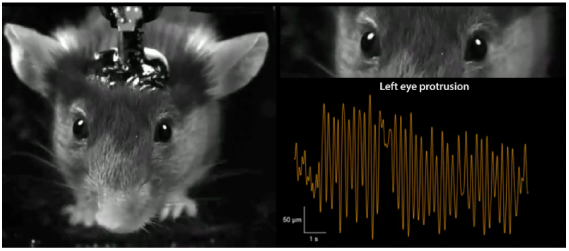
Example eye protrusion during chewing (slowed down 2*×*)

### Supplemental Tables

**Supplementary Table 1:**
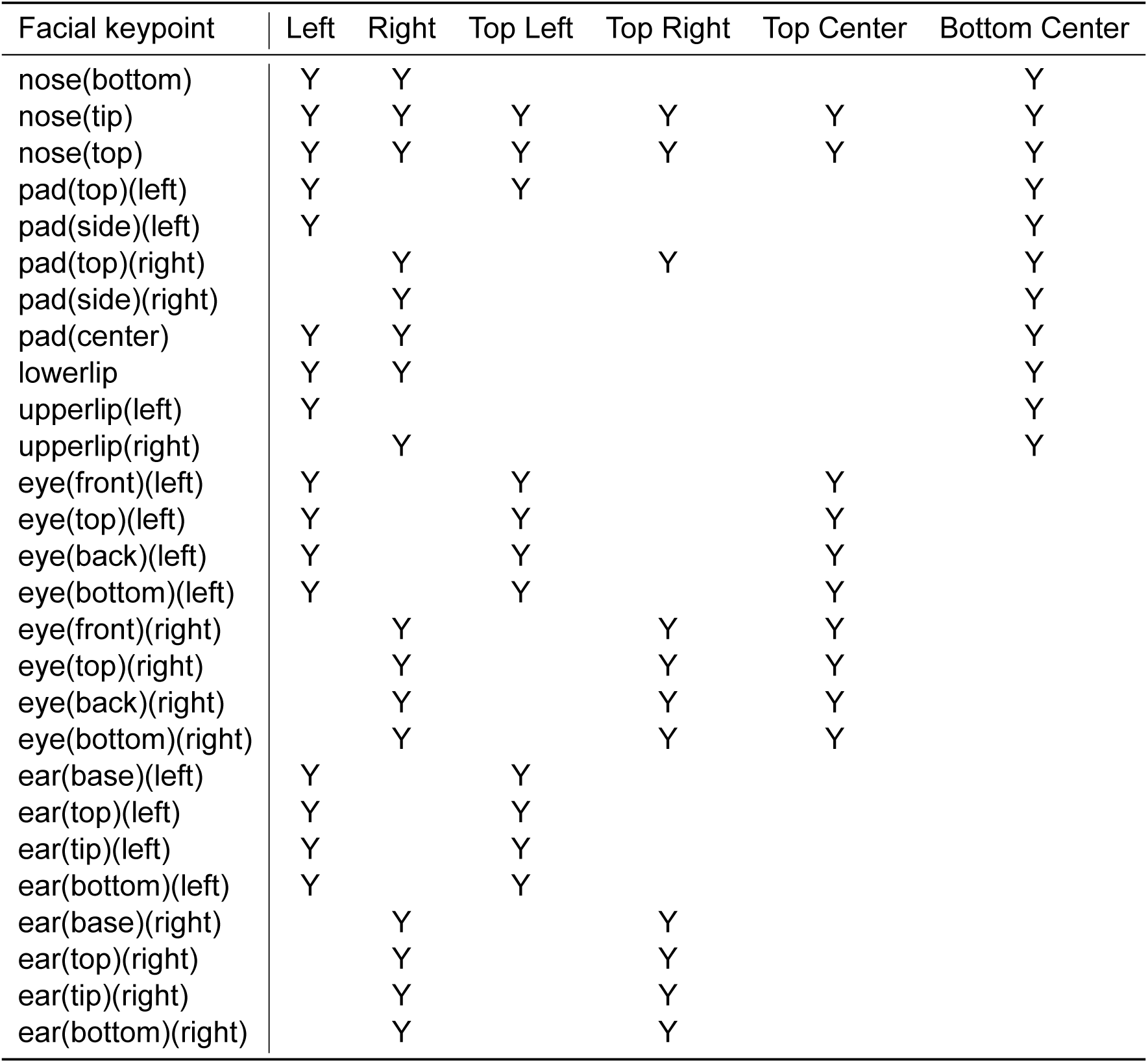
Keypoints labeled per camera view.

| Facial keypoint | Left | Right | Top Left | Top Right | Top Center | Bottom Center |
| --- | --- | --- | --- | --- | --- | --- |
| nose(bottom) | Y | Y |  |  |  | Y |
| nose(tip) | Y | Y | Y | Y | Y | Y |
| nose(top) | Y | Y | Y | Y | Y | Y |
| pad(top)(left) | Y |  | Y |  |  | Y |
| pad(side)(left) | Y |  |  |  |  | Y |
| pad(top)(right) |  | Y |  | Y |  | Y |
| pad(side)(right) |  | Y |  |  |  | Y |
| pad(center) | Y | Y |  |  |  | Y |
| lowerlip | Y | Y |  |  |  | Y |
| upperlip(left) | Y |  |  |  |  | Y |
| upperlip(right) |  | Y |  |  |  | Y |
| eye(front)(left) | Y |  | Y |  | Y |  |
| eye(top)(left) | Y |  | Y |  | Y |  |
| eye(back)(left) | Y |  | Y |  | Y |  |
| eye(bottom)(left) | Y |  | Y |  | Y |  |
| eye(front)(right) |  | Y |  | Y | Y |  |
| eye(top)(right) |  | Y |  | Y | Y |  |
| eye(back)(right) |  | Y |  | Y | Y |  |
| eye(bottom)(right) |  | Y |  | Y | Y |  |
| ear(base)(left) | Y |  | Y |  |  |  |
| ear(top)(left) | Y |  | Y |  |  |  |
| ear(tip)(left) | Y |  | Y |  |  |  |
| ear(bottom)(left) | Y |  | Y |  |  |  |
| ear(base)(right) |  | Y |  | Y |  |  |
| ear(top)(right) |  | Y |  | Y |  |  |
| ear(tip)(right) |  | Y |  | Y |  |  |
| ear(bottom)(right) |  | Y |  | Y |  |  |

**Supplementary Table 2:**
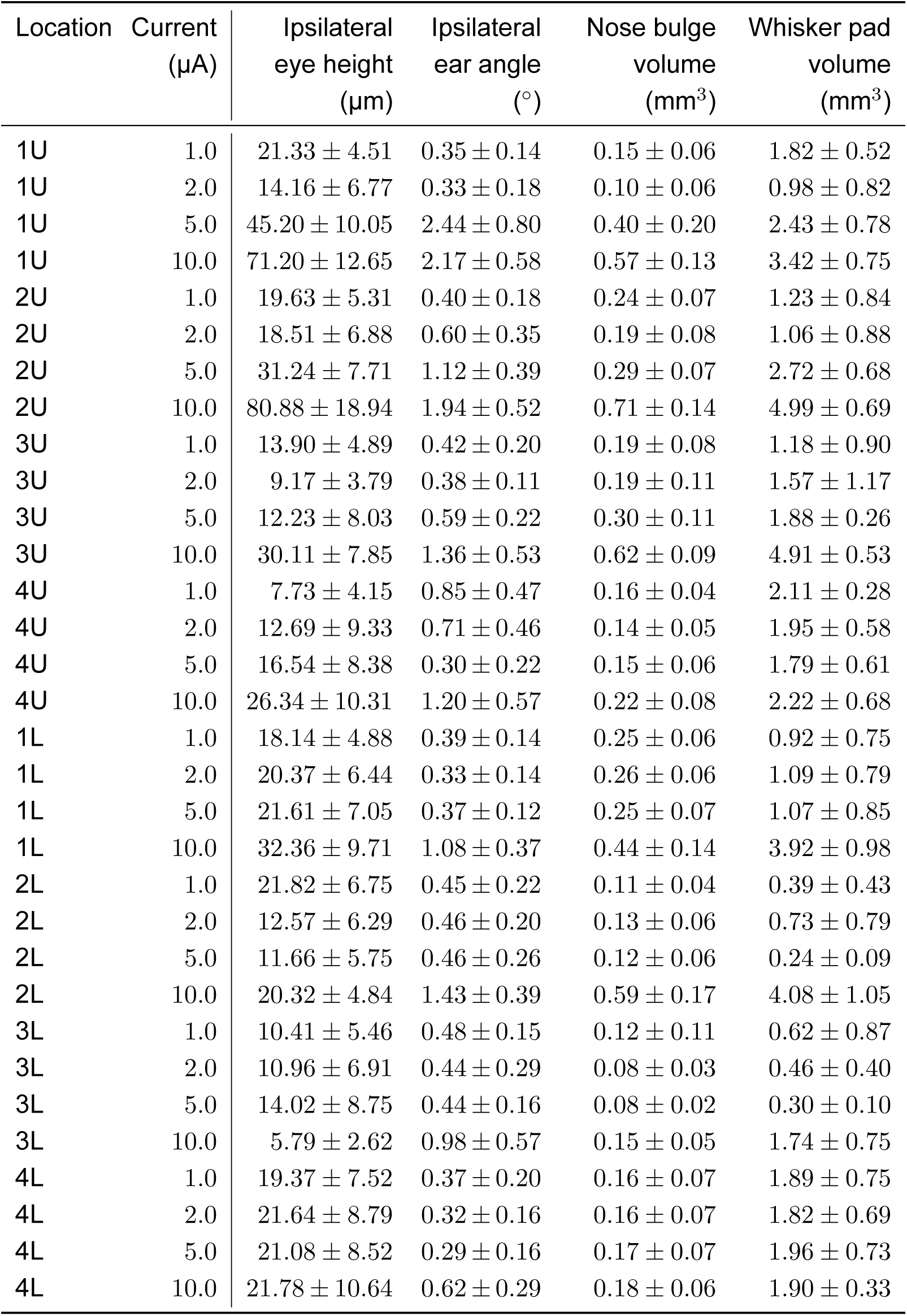
Summary statistics (mean *±* std) for Figure 5e.

